# Trim39 regulates neuronal apoptosis by acting as a SUMO-targeted E3 ubiquitin-ligase for the transcription factor NFATc3

**DOI:** 10.1101/2020.09.29.317958

**Authors:** Meenakshi Basu Shrivastava, Barbara Mojsa, Stéphan Mora, Ian Robbins, Guillaume Bossis, Iréna Lassot, Solange Desagher

**Affiliations:** IGMM, Univ Montpellier, CNRS, Montpellier, France; Centre for Gene Regulation and Expression, School of Life Science, University of Dundee, Dundee, UK

## Abstract

NFATc3 is the predominant member of the NFAT family of transcription factor in neurons, where it plays a pro-apoptotic role. Mechanisms controlling NFAT protein stability are poorly understood. Here we identify Trim39 as an E3 ubiquitin-ligase of NFATc3. Indeed, Trim39 ubiquitinates NFATc3 *in vitro* and in cells, whereas silencing of endogenous Trim39 decreases NFATc3 ubiquitination. We also show that Trim17 inhibits Trim39-mediated ubiquitination of NFATc3 by reducing both the E3 ubiquitin-ligase activity of Trim39 and the NFATc3/Trim39 interaction. Moreover, mutation of SUMOylation sites in NFATc3 or SUMO-interacting motif in Trim39 reduces the NFATc3/Trim39 interaction and Trim39-induced ubiquitination of NFATc3. As a consequence, silencing of Trim39 increases the protein level and transcriptional activity of NFATc3, resulting in enhanced neuronal apoptosis. Likewise, a SUMOylation-deficient mutant of NFATc3 exhibits increased stability and pro-apoptotic activity. Taken together, these data indicate that Trim39 modulates neuronal apoptosis by acting as a SUMO-targeted E3 ubiquitin-ligase for NFATc3.

## Introduction

The NFAT (Nuclear Factor of Activated T cells) family of transcription factors is a key player in a wide range of physiological and pathological processes. Initially discovered in activated T cells (Shaw et al., 1988), the different members of the NFAT family have been identified in most tissues where they play both redundant and specific roles (Fric et al., 2012; Kipanyula et al., 2016; Mognol et al., 2016; Wu et al., 2007). They are implicated in the development and the function of the immune system, brain, cardiovascular system, skeletal muscles, bones and other organs by regulating the expression of different target genes involved in cytokine production but also in cell proliferation, differentiation and apoptosis. As a consequence, NFAT deregulation is involved in many pathologies including auto-immune diseases, cancer and neurodegenerative diseases (Kipanyula et al., 2016; J.-U. Lee et al., 2018; Müller & Rao, 2010). A better understanding of NFAT regulation, in particular by post-translational modification and degradation, is therefore of crucial importance.

The calcium-regulated, cytoplasmic-nuclear shuttling of NFATc1, NFATc2, NFATc3 and NFATc4 has been extensively studied. These NFAT members are normally found in the cytoplasm in a hyperphosphorylated and inactive state. Upon an increase in intracellular calcium levels, they are dephosphorylated by the calcium-dependent phosphatase calcineurin, which triggers their nuclear import and activation. Once in the nucleus, NFATs induce (or repress) the transcription of specific target genes, usually in cooperation with partner transcription factors such as AP-1 or co-activators (Hogan et al., 2003; Mognol et al., 2016; Müller & Rao, 2010). In contrast, the regulation of NFAT stability by the ubiquitin-proteasome system remains elusive. Indeed, only a few studies have addressed this issue. However, NFATs are relatively short-lived proteins and previous studies have shown that interfering with the regulation of NFAT levels by the ubiquitin-proteasome system can have a marked impact on the physiology of various cell types (Chao et al., 2019; X. Li et al., 2015; Narahara et al., 2019; Singh et al., 2011; Yoeli-Lerner et al., 2005; Youn et al., 2012). In addition to phosphorylation and ubiquitination, NFAT proteins have been shown to be regulated by SUMOylation. Several studies have shown that covalent conjugation of SUMO to NFATs has an impact on their cytoplasmic-nuclear shuttling, subnuclear localization and transcriptional activity (E. T. Kim et al., 2019; Nayak et al., 2009; Terui et al., 2004; Vihma & Timmusk, 2017). Indeed, SUMOylation can have many consequences on its substrate proteins, including modification of their activity, interaction properties and subcellular localization (Henley et al., 2018; X. Zhao, 2018). In addition, SUMOylation of proteins can regulate their stability (Liebelt & Vertegaal, 2016). Indeed, a few E3 ubiquitin-ligases that specifically recognize and ubiquitinate SUMOylated proteins have been described (Geoffroy & Hay, 2009; Prudden et al., 2007; Sriramachandran & Dohmen, 2014). These SUMO-targeted E3 ubiquitin-ligases (STUbLs) generally induce the degradation of their substrates by the proteasome, raising the possibility that SUMO might also modulate NFAT ubiquitination and degradation.

NFATc3 is the predominant NFAT family member expressed in various neuronal types (M. S. Kim & Usachev, 2009; Luo et al., 2014; Mojsa et al., 2015; Ulrich et al., 2012; Vashishta et al., 2009). We have previously shown that NFATc3 is involved in the regulation of neuronal apoptosis (Mojsa et al., 2015). Two independent studies have also implicated NFATc3 in α-synuclein-induced degeneration of midbrain dopaminergic neurons in Parkinson’s disease (Caraveo et al., 2014; Luo et al., 2014). Interestingly, following depolarization-induced elevations of intracellular calcium concentration in neurons, NFATc3 is more rapidly and strongly activated than NFATc4, (Ulrich et al., 2012). Once in the nucleus, activation of pro-apoptotic protein kinases such as GSK3β does not seem to be sufficient to induce NFATc3 nuclear exclusion in neurons (Mojsa et al., 2015; Ulrich et al., 2012). Proteasomal degradation could therefore be an alternative way to reduce its activity in this case. However, only one study relating NFATc3 ubiquitination and degradation has been reported so far and this in the context of LPS-induced cardiac hypertrophy (Chao et al., 2019). In previous work, we have shown that NFATc3 can be SUMOylated on three consensus sites (Mojsa et al., 2015). We have also found that NFATc3 binds to Trim17 (Mojsa et al., 2015), which belongs to a large family of RING-containing E3 ubiquitin-ligases. Although its E3 ubiquitin-ligase activity has been confirmed (I. Lassot et al., 2010; Urano et al., 2009), Trim17 does not induce NFATc3 ubiquitination. On the contrary, overexpression of Trim17 reduces the ubiquitination of NFATc3 and increases its steady-state protein level (Mojsa et al., 2015). Since TRIM17 can prevent ubiquitination of some of its binding partners by inhibiting other E3 ubiquitin-ligases from the TRIM family (Iréna Lassot et al., 2018; Lionnard et al., 2019), we hypothesized that the stability of NFATc3 might be regulated by a TRIM protein interacting with Trim17, such as Trim39.

In the present study, we demonstrate that Trim39 is a genuine E3 ubiquitin-ligase for NFATc3. We also show that Trim39-mediated ubiquitination of NFATc3 is inhibited by Trim17. Moreover, mutation of NFATc3 SUMOylation sites both decreases its ubiquitination by Trim39 and increases its stability. The same effects are reproduced by mutation of a crucial SUMO-interacting motif (SIM) in Trim39. These data indicate that Trim39 acts as a STUbL for NFATc3. As a result, SUMO and Trim39 modulate the transcriptional activity of NFATc3 and its pro-apoptotic effect in neurons. Therefore, our study provides the identification of a new STUbL and a first insight into complex mechanisms regulating the stability of NFATc3 in neurons.

## Results

### Trim39 is an E3 ubiquitin-ligase for NFATc3

Human TRIM39 and TRIM17 proteins have been found to interact with each other in three independent proteome-scale yeast two-hybrid screens (Rolland et al., 2014; Rual et al., 2005; Woodsmith et al., 2012). To determine whether mouse Trim39 and Trim17 proteins can also bind to each other, and whether Trim39 can bind to NFTAc3, co-immunoprecipitation experiments were performed. Indeed, in cells co-transfected with Trim17-GFP and Flag-Trim39, immunoprecipitation of Trim39 using anti-Flag antibody co-precipitated Trim17, whereas immunoprecipitation of Trim17 using GFP-Trap beads co-precipitated Trim39 (Fig. 1A). In a similar way, in cells co-transfected with HA-NFATc3 and Flag-Trim39, the two proteins were reciprocally co-immunoprecipitated by using either anti-Flag or anti-HA antibodies (Fig. 1B). To confirm this interaction at the endogenous level, we next performed an *in situ* proximity ligation assay (PLA) in Neuro2A cells, using anti-NFATc3 and anti-Trim39 antibodies. Close proximity was detected between endogenous NFATc3 and endogenous Trim39 as assessed by a PLA signal (Fig. 1C) that was predominantly cytoplasmic (Fig. 1C, endo 1 slice). Overexpression of Trim39 increased the PLA signal (Fig. 1C). To confirm the specificity of the assay, we used a specific shRNA against Trim39 and first verified that it effectively reduces the level of endogenous Trim39 protein (Fig. 1D). As expected, silencing of Trim39 using this shRNA strongly decreased the PLA signal (Fig. 1E). Taken together, these data indicate that Trim39 interacts with both Trim17 and NFATc3.

**Figure 1.**
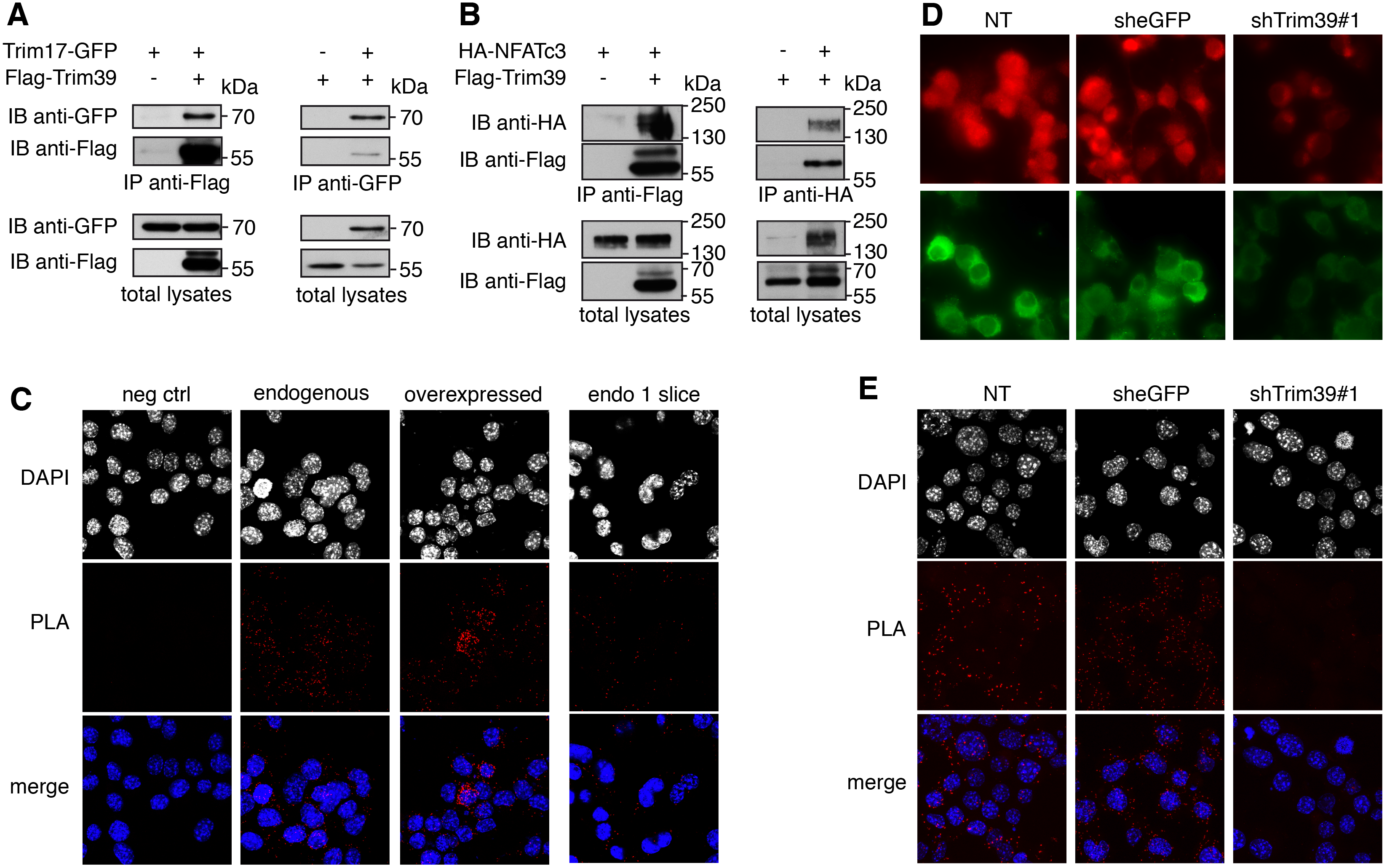
Trim39 interacts with both Trim17 and NFATc3. **A.** BHK cells were transfected with Trim17-GFP together with Flag-Trim39 or empty plasmid (as a negative control) for 24 h. Cells were then treated with 20 μM MG-132 for 5 h. The cells were subsequently harvested and lysates were subjected to immunoprecipitation using anti-Flag agarose beads (left panel) or GFP-Trap beads (right panel). Immunoprecipitates and total lysates were analyzed by western blot using anti-GFP and anti-Flag antibodies. **B.** Neuro2A cells were transfected with HA-NFATc3 together with Flag-Trim39 or empty plasmid for 24 h. Cells were then treated as in A and lysates were subjected to immunoprecipitation using anti-Flag (left panel) or anti-HA (right panel) antibodies. Immunoprecipitates and total lysates were analyzed by western blot using anti-HA and anti-Flag antibodies. **C.** Neuro2A cells were treated with 10 μM MG-132 for 4 h and then fixed and subjected to *in situ* PLA using rabbit anti-NFATc3 and mouse anti-Trim39 antibodies. Each bright red spot indicates that the two proteins are in close proximity. Negative control was obtained by omitting anti-Trim39 antibody. When indicated, cells were previously transfected with Trim39 for 24 h (overexpressed). Images were analyzed by confocal microscopy. To better visualize the differences in PLA intensity, maximum intensity projection was applied to the z-stacks of images on the left panel. To better determine the subcellular location of the NFATc3/Trim39 interaction, a single slice of the z-stack is presented on the right panel (endo 1 slice). Nuclear staining was performed using DAPI. **D.** Neuro2A cells were transduced with lentiviral particles expressing a control shRNA (sheGFP) or a specific shRNA against Trim39 (shTrim39#1) for 24 h. Transduced cells were selected using puromycin for two additional days and plated onto coverslips. The day after plating, cells were analyzed by immunofluorescence using two different antibodies against Trim39; in red: antibody from Origene, in green: antibody from Proteintech. Images were set to the same minimum and maximum intensity to allow signal intensity comparison. **E.** Additional coverslips from the experiment presented in D were treated as in C and z-stacks of images were subjected to maximum intensity projection.

We next examined whether Trim39 could mediate the ubiquitination of NFATc3. In cells co-transfected with His-tagged ubiquitin, the ubiquitination level of NFATc3 was significantly increased by overexpressed Trim39 but not by an inactive mutant deleted of its RING domain (Trim39-ΔRING; Fig. 2A). In contrast, silencing of Trim39 using three different specific shRNAs, deeply decreased the ubiquitination of NFATc3 (Fig. 2B), indicating that endogenous Trim39 is involved in the ubiquitination of NFATc3. To further demonstrate that Trim39 is an E3 ubiquitin-ligase of NFATc3, we carried out an *in vitro* ubiquitination assay using *in vitro* translated/immuno-purified NFATc3 and purified recombinant proteins. In these experiments, GST-Trim39 stimulated NFATc3 ubiquitination in the presence of ubiquitin, E1 and E2 enzymes but not in the absence of ubiquitin (Fig. 2C). In contrast, an inactive mutant of Trim39 in which two crucial Cys residues of the RING domain where mutated (GST-Trim39-C49S/C52S) did not have any effect (Fig. 2C). Taken together, these data indicate that NFATc3 is a direct substrate for the E3 ubiquitin-ligase activity of Trim39.

**Figure 2.**
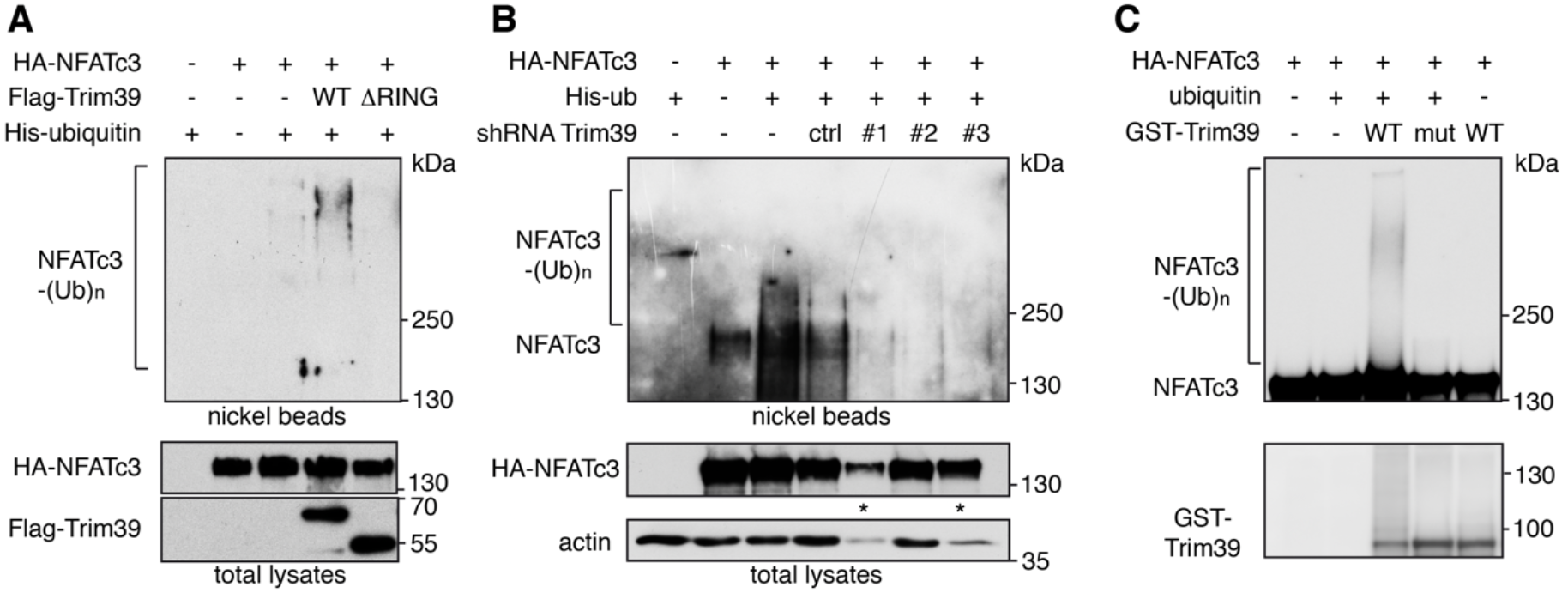
Trim39 is an E3 ubiquitin-ligase of NFATc3. **A.** Neuro2A cells were transfected with HA-NFATc3 together with empty plasmid or His-tagged ubiquitin, in the presence or absence of Flag-Trim39 or the inactive mutant Flag-Trim39-ΔRING for 24 h. Cells were then incubated with 20 μM MG-132 for 6 h before harvesting. The ubiquitinated proteins were purified using nickel beads and analyzed by western blotting using anti-HA antibody to detect ubiquitin-conjugated HA-NFATc3. In a separate SDS-PAGE, samples of the input lysates used for the purification were analyzed with antibodies against HA and Flag. **B.** Neuro2A cells were transduced with lentiviruses expressing a control shRNA (directed against eGFP) or three different shRNAs targeting Trim39. Following 24 h transduction and 48 h selection of transduced cells using puromycin, cells were plated in new dishes. Then, cells were transfected with HA-NFATc3 or His-tagged ubiquitin or both for 24 h, and treated as in A. In conditions indicated by a star (*), some material was lost during TCA precipitation of the input lysate without affecting the amount of proteins in the nickel bead purification. These data are representative of 4 independent experiments. **C.** *In vitro* translated HA-NFATc3 was first immunopurified from wheat germ extract using anti-HA antibody. Then, beads used for immunopurification of NFATc3 were incubated for 1 h at 37°C in the *in vitro* ubiquitination reaction mix (containing E1 and E2 enzymes) with purified recombinant GST-Trim39 (WT) or its inactive mutant GST-Trim39-C49S/C52S (mut) in the presence or the absence of ubiquitin as indicated. Poly-ubiquitinated forms of NFATc3 were detected by immunoblotting using an anti-NFATc3 antibody. The same membrane was immunoblotted with an anti-TRIM39 antibody to verify that similar amounts of recombinant WT GST-Trim39 and GST-Trim39-C49S/C52S were used in the assay. Note that in the presence of ubiquitin the unmodified form of WT GST-Trim39 is lower due to high Trim39 ubiquitination.

### Trim39 induces the degradation of NFATc3 and decreases its transcriptional activity

Because ubiquitination often targets proteins for proteasomal degradation, we examined whether Trim39 could impact the protein level of NFATc3. Indeed, the level of NFATc3 progressively decreased when co-transfected with increasing amounts of Trim39 (Fig. 3A). Interestingly, the inactive mutant Trim39-ΔRING did not decrease the protein level of NFATc3 but rather increased it, in a similar way as the proteasome inhibitor MG-132 (Fig. 3A). Mutations of the RING domain of E3 ubiquitin-ligases generally induce a dominant-negative effect (I. Lassot et al., 2010; Pickart, 2001). Therefore, this increase in NFATc3 protein may be due to the inhibition of endogenous Trim39 by Trim39-ΔRING, as it has been previously reported for the effect of Trim39 on the half-life of p53 (Zhang, Huang, et al., 2012). Consistently, silencing of endogenous Trim39 using a specific siRNA also significantly increased the protein level of endogenous NFATc3 in Neuro2A cells (Fig. 3B). Taken together, these data strongly suggest that Trim39-mediated ubiquitination is involved in the proteasomal degradation of NFATc3.

**Figure 3.**
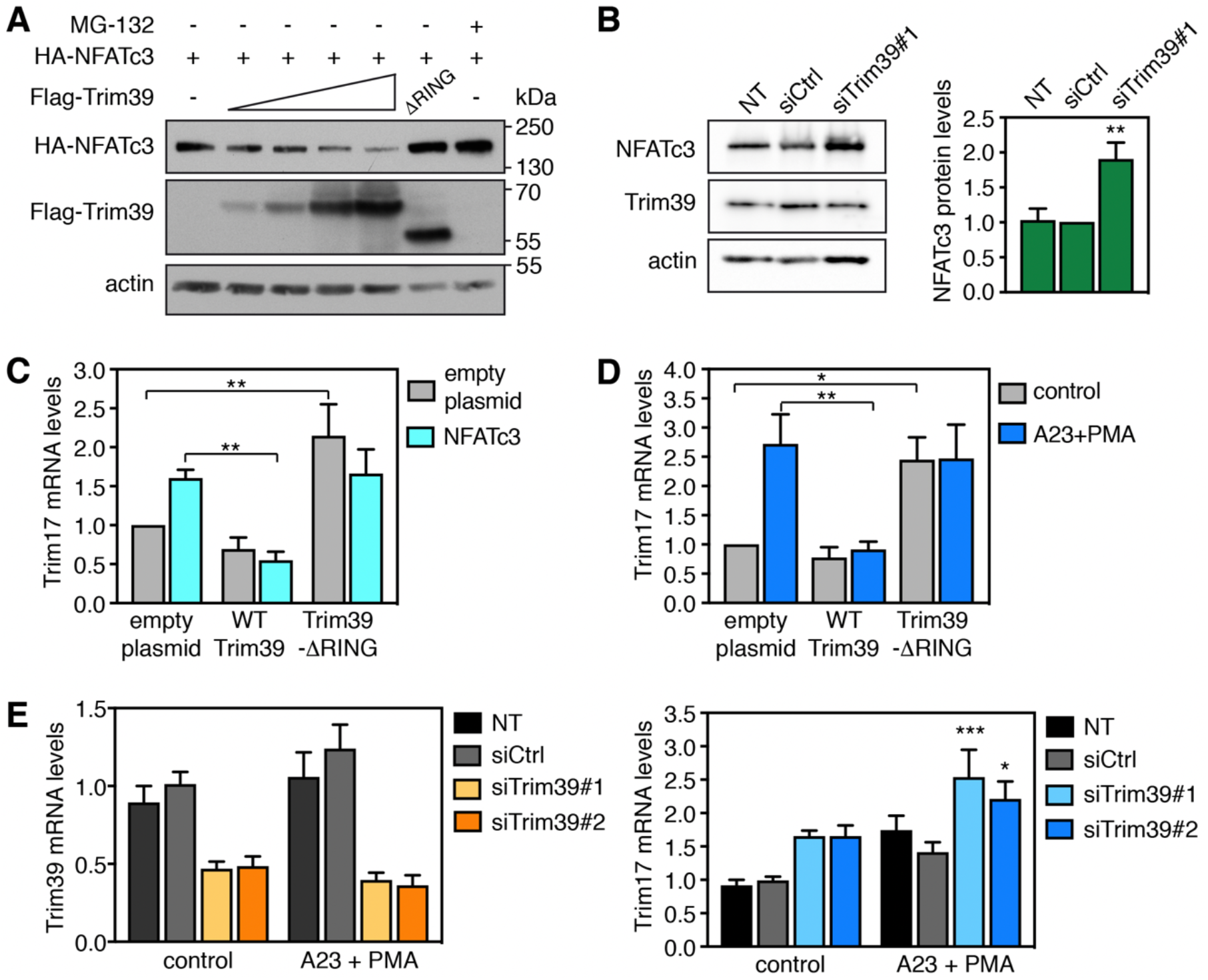
Trim39 mediates NFATc3 degradation. **A.** BHK cells were transfected with a fixed amount of a HA-NFATc3 expressing vector (1 μg) together with empty plasmid (−) or increasing amounts of Flag-Trim39 expressing vector (0.1, 0.2, 0.5 and 1 μg) or 0.2 μg of a vector expressing the inactive mutant Flag-Trim39-ΔRING. When indicated, the cells were treated with 10 μM MG-132 for 6 h before harvesting. Total lysates were analyzed by western blot using antibodies against HA, Flag and actin. **B.** Neuro2A cells were left untreated (NT), or transfected twice with an siRNA targeting specifically Trim39 (siTrim39#1) or with a negative control siRNA (siCtrl) for 48 h. Total lysates were analyzed by western blot using antibodies against NFATc3, Trim39 and actin. The intensity of the NFATc3 bands on the western blots was quantified, normalized by the intensity of the actin bands and expressed relative to the values obtained with the control shRNA. The graph shows mean ± SEM from three independent experiments. **P<0.01 significantly different from siCtrl (one-way ANOVA followed by Dunnett’s multiple comparisons test). **C.** Neuro2A cells were co-transfected with empty plasmid or HA-NFATc3, together with empty plasmid, Flag-Trim39 or the inactive mutant Flag-Trim39-ΔRING for 24 h. Then, total RNA was extracted and the mRNA level of Trim17 was estimated by quantitative PCR. Data are the means ± SEM of four independent experiments. **P<0.01 significantly different from the corresponding control (two-way ANOVA followed by Sidak’s multiple comparisons test). **D.** Neuro2A cells were transfected with empty plasmid, Flag-Trim39 or Flag-Trim39-ΔRING for 24 h. Then, cells were left untreated (control) or were deprived of serum for 3 h and subsequently treated with 1 μM A23187 and 100 nM PMA in serum-free medium for 1 h (A23+PMA). Total RNA was extracted and the mRNA level of Trim17 was estimated by quantitative PCR. Data are the means ± SEM of three independent experiments. *P<0.05; **P<0.01 significantly different from the corresponding value in cells transfected with empty plasmid (two-way ANOVA followed by Sidak’s multiple comparisons test). **E.** Neuro2A cells were transfected twice with two different siRNAs targeting specifically Trim39 or with a negative control siRNA for 48 h. Then, cells were left untreated (control) or were deprived of serum for 3h and subsequently treated with 1 μM A23187 and 100 nM PMA in serum-free medium for 30 min (A23+PMA). Total RNA was extracted and the mRNA level of Trim39 (left panel) or Trim17 (right panel) was estimated by quantitative PCR (NT = non transfected). Data are the means ± SEM of six independent experiments. *P<0.05; ***P<0.001 significantly different from cells transfected with control siRNA in the same condition (two-way ANOVA followed by Sidak’s multiple comparisons test).

To examine whether the effect of Trim39 on the protein level of NFATc3 could have an impact on its activity as a transcription factor, we measured the mRNA level of one of its target genes: *Trim17*. Indeed, in a previous study, we have shown that Trim17 is transcriptionally induced by NFATc3 (Mojsa et al., 2015). Consistently, in the present study, Trim17 mRNA level was increased when NFATc3 was overexpressed (Fig. 3C). Interestingly, this induction was completely abrogated by co-expression of wild type but not inactive Trim39 (ΔRING). Moreover, even when NFATc3 was not transfected, Trim39-ΔRING significantly increased the expression level of Trim17 (Fig. 3C), suggesting that the inhibition of endogenous Trim39 through a dominant negative effect is sufficient to increase the activity of endogenous NFATc3 (Fig. 3A). To confirm these data, Neuro2A cells were treated with the calcium ionophore A23187 and phorbol 12-myristate 13-acetate (PMA) to activate both endogenous NFATc3 (through calcium-induced nuclear translocation) and its transcriptional partner AP-1. As previously reported (Mojsa et al., 2015), Trim17 mRNA level was increased following treatment with A23187 and PMA (Fig. 3D). Although increase in intracellular calcium should activate other members of the NFAT family, this induction of Trim17 is probably due to NFATc3 as it is the NFAT transcription factor that is predominantly expressed in Neuro2A cells (Mojsa et al., 2015). Again, this Trim17 induction was completely abrogated by overexpression of wild type Trim39 but not inactive Trim39 (Fig. 3D). Overexpression of the dominant-negative mutant Trim39-ΔRING also significantly increased the expression level of Trim17, even in control conditions (Fig. 3D). Taken together, these data suggest that exogenous Trim39 reduces the transcriptional activity of both overexpressed and endogenous NFATc3. To determine the impact of endogenous Trim39, Neuro2A cells were transfected with two different specific siRNAs that efficiently decreased the mRNA level of Trim39 (Fig. 3E, left panel). Silencing of Trim39 resulted in Trim17 induction, notably following treatment with A23187 and PMA, which activates endogenous NFATc3 (Fig. 3E, right panel). These data therefore indicate that endogenous Trim39 also regulates endogenous NFATc3. As we have previously shown that Trim17 can bind and inhibit NFATc3 by preventing its nuclear translocation (Mojsa et al., 2015), we examined whether Trim39 could have the same effect on NFATc3. Indeed, under conditions where Trim17 decreased the nuclear translocation of NFATc3 by more than twofold, Trim39 had no impact on the subcellular localization of NFATc3 (Fig. S1). Therefore, these data strongly suggest that Trim39 inhibits the transcription factor activity of NFATc3 by ubiquitinating it and by inducing its proteasomal degradation, but not by preventing its nuclear translocation.

### Trim17 inhibits the ubiquitination of NFATc3 mediated by Trim39

As we initially observed that Trim17 decreases the ubiquitination level of NFATc3 (Mojsa et al., 2015), we tested whether Trim17 could affect Trim39-mediated ubiquitination of NFATc3. Indeed, the increase in NFATc3 ubiquitination induced by Trim39 overexpression was abolished by the co-expression of Trim17 in cells (Fig. 4A). This effect was confirmed *in vitro*. Indeed, the ubiquitination of *in vitro* translated NFATc3 by recombinant His-TRIM39 was completely prevented by recombinant MBP-TRIM17 (Fig. 4B). As only purified proteins were used in a complete acellular medium for this assay, these results suggest that TRIM17 can directly inhibit the ubiquitination of NFATc3 induced by TRIM39. Interestingly, the ubiquitination level of Trim39 was strongly decreased in the presence of Trim17 both in cells (Fig. 4A) and *in vitro* (Fig. 4B), excluding the possibility that Trim17 acts by ubiquitinating Trim39. Moreover, as *in vitro* auto-ubiquitination gives a measure of E3 ubiquitin-ligase activity (Pickart, 2001), this also suggests that TRIM17 can directly inhibit the E3 ubiquitin-ligase activity of TRIM39. Interestingly, in these experiments, the *in vitro* auto-ubiquitination of TRIM17 was also decreased (Fig. 4B) by TRIM39 and the ubiquitination level of Trim17 in cells was also reduced in the presence of Trim39 (Fig. S2), suggesting a reciprocal inhibition of the two TRIM proteins.

**Figure 4.**
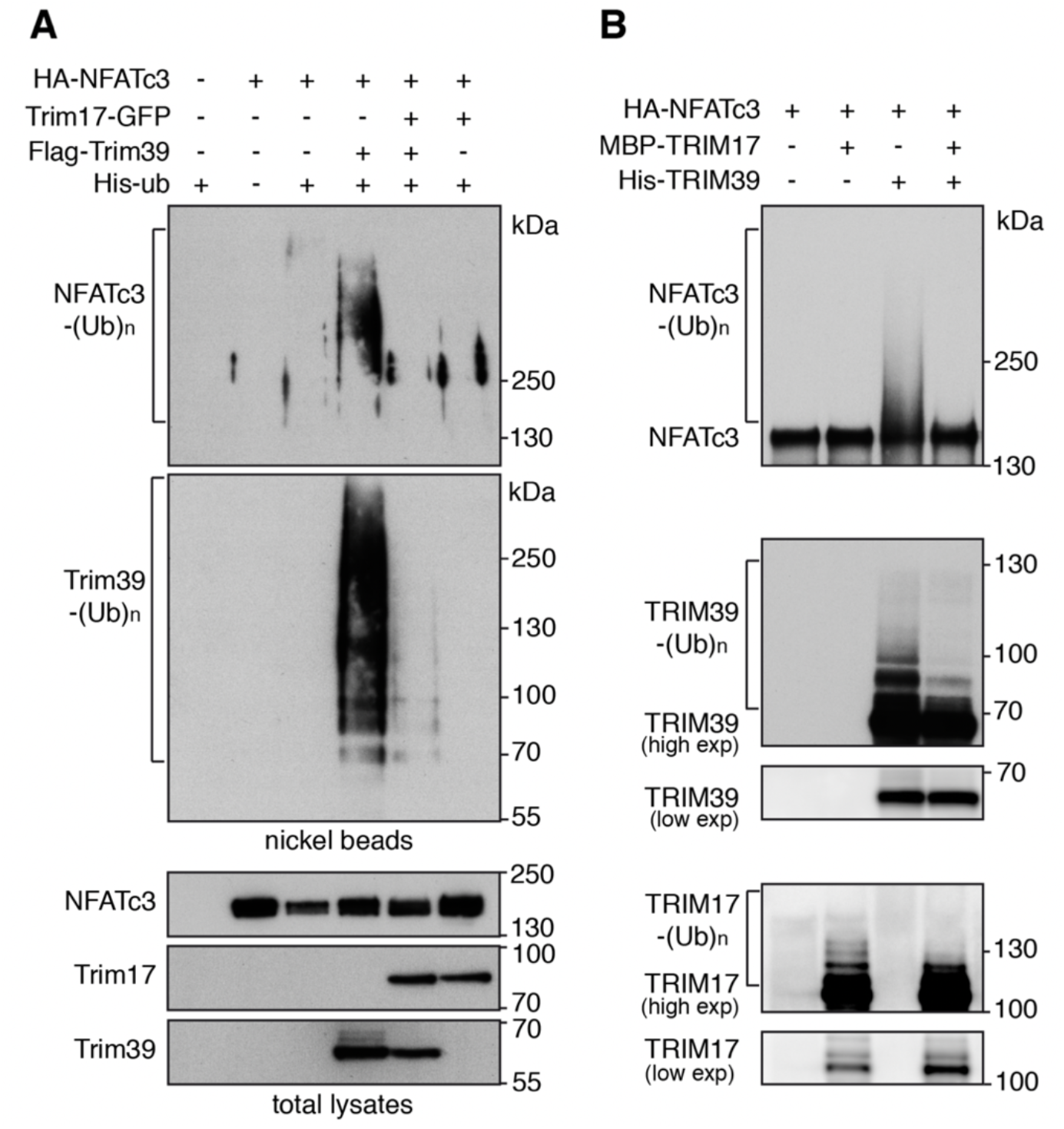
Trim17 inhibits TRIM39-mediated uiquitination of NFATc3. **A.** BHK cells were transfected with HA-NFATc3 together with His-tagged ubiquitin, in the presence or the absence of Flag-Trim39, Trim17-GFP or both, as indicated, for 24 h. Then, cells were incubated with 20 μM MG-132 for 6 h before harvesting. The ubiquitinated proteins were purified using nickel beads and analyzed by western blotting using anti-HA and anti-Flag antibodies to detect poly-ubiquitinated forms of NFATc3 and Trim39. In a separate SDS-PAGE, samples of the input lysates used for the purification were analyzed with antibodies against HA, Flag and GFP. **B.** *In vitro* translated HA-NFATc3 was first immunopurified from wheat germ extract using anti-HA antibody. Then, beads used for immunopurification of NFATc3 were incubated for 1 h at 37°C in the *in vitro* ubiquitination reaction mix (containing ubiquitin and E1 and E2 enzymes) with purified recombinant His-TRIM39 or MBP-TRIM17 as indicated. Poly-ubiquitinated forms of NFATc3, TRIM39 and TRIM17 were detected by immunoblotting using anti-NFATc3, anti-TRIM39 and anti-TRIM17 antibodies revealed using high exposure times. Low exposure times were used to compare the level of TRIM39 and TRIM17 in the different conditions.

To further investigate the mechanisms underlying the inhibitory effect of Trim17, the impact of Trim17 on the interaction between NFATc3 and Trim39 was assessed. Strikingly, when co-transfected with HA-NFATc3 and Flag-Trim39, Trim17-GFP almost completely prevented the co-immunoprecipitation of Flag-Trim39 with HA-NFATc3 (Fig. 5A) or the co-immunoprecipitation of HA-NFATc3 with Flag-Trim39 (Fig. 5B). Moreover, in PLA experiments, the close proximity signal between endogenous NFATc3 and Trim39 proteins was significantly reduced by the overexpression of Trim17-GFP compared to GFP (Fig. 5C,D).

**Figure 5.**
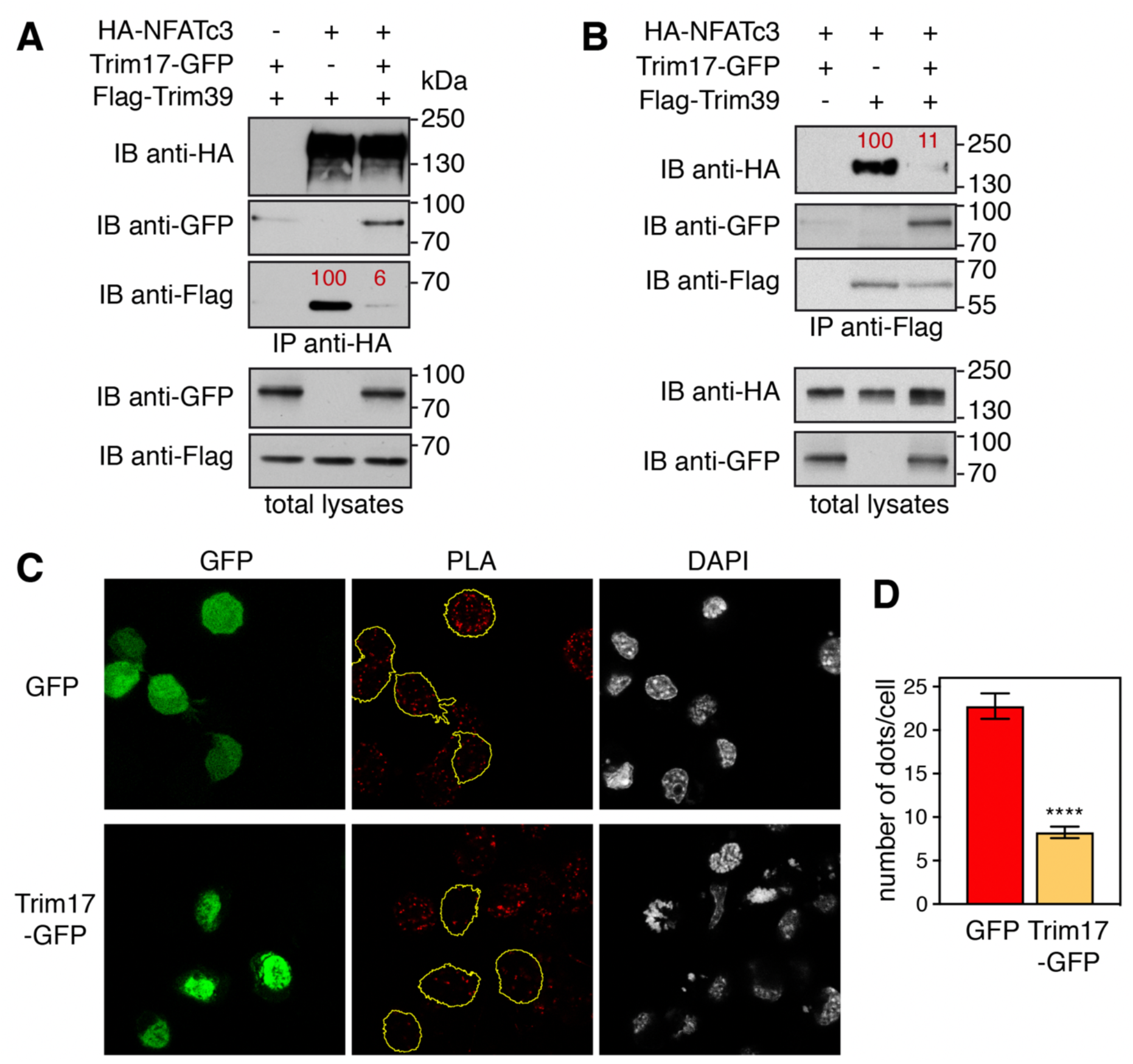
Trim17 reduces the interaction between endogenous Trim39 and NFATc3. **A,B.** Neuro2A cells were transfected with HA-NFATc3 in the presence or the absence of Flag-Trim39, Trim17-GFP or both, as indicated, for 24 h. Cells were then treated with 20 μM MG-132 for 7 h. The cells were subsequently harvested and lysates were subjected to immunoprecipitation using anti-HA (A) or anti-Flag (B) antibodies. Immunoprecipitates and total lysates were analyzed by western blot using anti-HA, anti-GFP and anti-Flag antibodies. The intensity of the bands containing Flag-Trim39 co-immunoprecipitated with HA-NFATc3 was normalized by the intensity of the bands corresponding to immunoprecipitated HA-NFATc3 (A). The intensity of the bands containing HA-NFATc3 co-immunoprecipitated with Flag-Trim39 was normalized by the intensity of the bands corresponding to immunoprecipitated Flag-Trim39 (B). Relative values are indicated in red. **C.** Neuro2A cells were transfected with GFP or Trim17-GFP for 24 h. Then cells were treated with 10 μM MG-132 for 4 h, fixed and subjected to *in situ* PLA using rabbit anti-NFATc3 and mouse anti-Trim39 antibodies. Each bright red spot indicates that the two proteins are in close proximity. Images were analyzed by confocal microscopy and a single slice of the z-stacks is presented for each condition. Nuclear staining was performed using DAPI. Note that, in the Trim17-GFP condition, transfected cells (delineated by a yellow line) show less dots than neighboring non transfected cells, which is not the case in the GFP condition. **D.** The number of dots was determined in individual cells transfected with either GFP or Trim17-GFP using Fiji. Data represent one experiment, including 68 transfected cells for each condition, representative of two independent experiments. ****p<0.0001, significantly different from GFP transfected cells (unpaired t test).

Taken together, these data strongly suggest that Trim17 inhibits the ubiquitination of NFATc3 mediated by Trim39 by inhibiting both the intrinsic E3 ubiquitin-ligase activity of Trim39 and the interaction between NFATc3 and Trim39.

### SUMOylation of NFATc3 modulates its ubiquitination and stability

In a previous study, we have identified three consensus SUMOylation sites in NFATc3 (Mojsa et al., 2015). As SUMOylation can modify the stability of proteins (Liebelt & Vertegaal, 2016), we tested whether alteration of the SUMOylation of NFATc3 can have an impact on its ubiquitination and half-life. We had previously used NFATc3 K/R mutants in which the acceptor Lys residues of the SUMOylation consensus motifs were replaced by Arg (Mojsa et al., 2015). However, large-scale mass spectrometry studies have shown that a quarter of SUMO acceptors lysines are also used for ubiquitin modification (Liebelt & Vertegaal, 2016). Therefore, additional NFATc3 mutants were generated in order to prevent SUMOylation without affecting a possible ubiquitination at these sites. For this purpose, the Glu residues of the NFATc3 SUMOylation consensus motifs (ΨK*X*E with Ψ representing a large hydrophobic residue and *X* any amino acid (Pichler et al., 2017; Rodriguez et al., 2001)) were substituted for Ala to generate NFATc3 E/A mutants. As expected, *in vitro* SUMOylation of the NFATc3-EallA mutant (in which the Glu residues of the three consensus motifs were replaced by Ala) and the NFATc3-KallR mutant (in which the Lys residues of the three consensus motifs were replaced by Arg), was almost completely abrogated (Fig. 6A). Interestingly, the ubiquitination level of NFATc3 in Neuro2A cells was not significantly altered by single or double E/A mutations whereas it was strongly decreased by the triple mutation (Fig. 6B), suggesting that SUMOylation of at least one consensus motif is necessary to favour the ubiquitination of NFATc3. Consistently, the half-life of the NFATc3-EallA mutant, measured after inhibition of protein synthesis with cycloheximide, was significantly increased compared to WT NFATc3 (Fig. 6C,D). Taken as a whole, these data suggest that SUMOylation of NFATc3 favours its ubiquitination and subsequent degradation.

**Figure 6.**
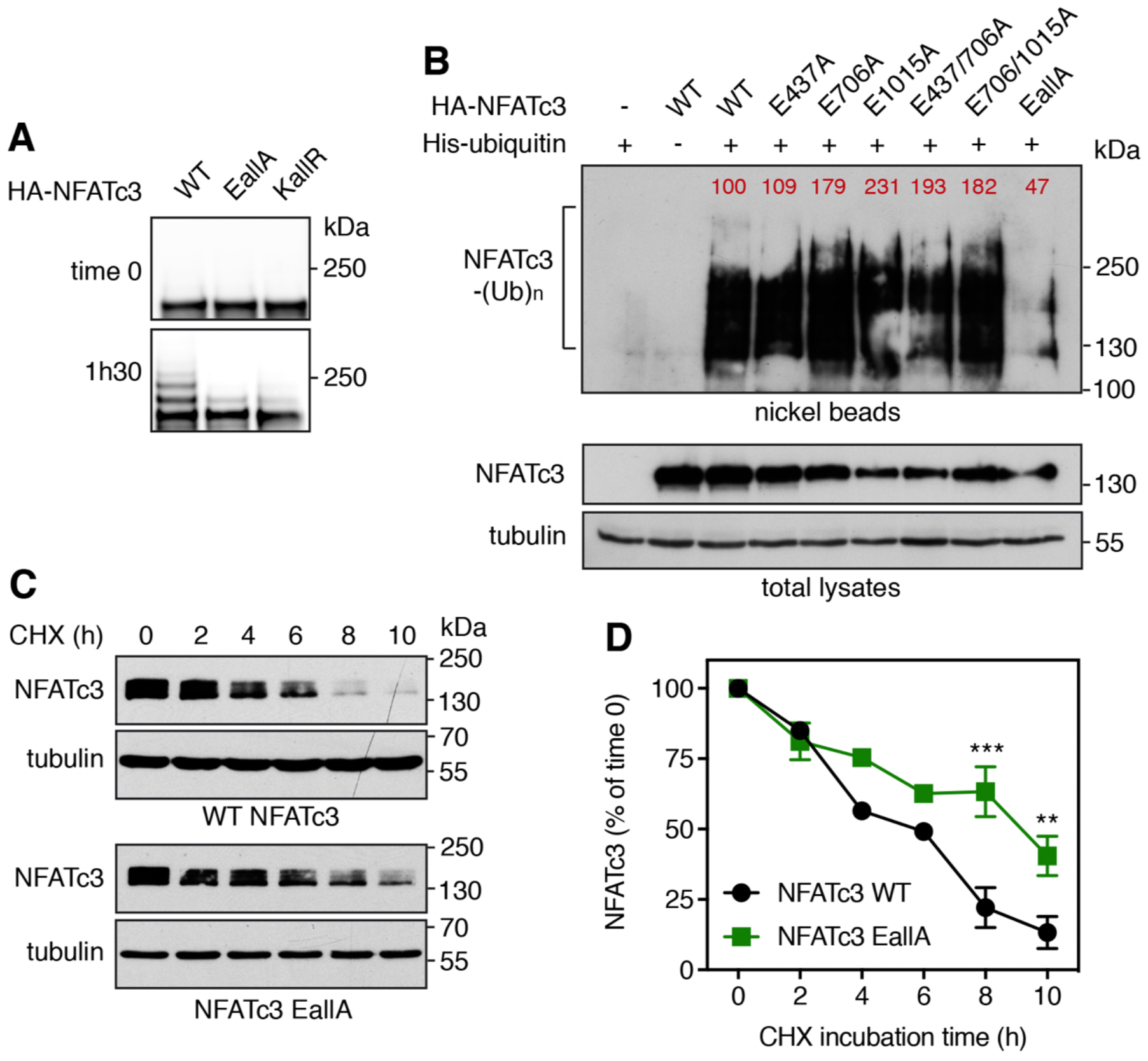
SUMOylation of NFATc3 modulates its ubiquitination and stability. **A.** *In vitro* translated HA-NFATc3 was incubated with *in vitro* SUMOylation reaction mix (containing SUMO1, E1, E2 and E3 enzymes) for 1 h 30 or directly added to sample loading buffer together with reaction mix (time 0). Multi-SUMOylated forms of NFATc3 were detected by immunoblotting using anti-NFATc3 antibody. **B.** Neuro2A cells were transfected with His-tagged ubiquitin or empty plasmid, together with WT HA-NFATc3 or the different HA-NFATc3 E/A mutant constructs for 24 h. Then, cells were incubated with 20 μM MG-132 for 6 h before harvesting. The ubiquitinated proteins were purified using nickel beads and analyzed by western blotting using anti-HA antibody to detect ubiquitin-conjugated HA-NFATc3. In a separate SDS-PAGE, samples of the input lysates used for the purification were analyzed with antibodies against HA and tubulin. The intensity of the NFATc3 bands from the nickel bead purification was normalized by the intensity of the bands in the total lysates. Relative values are indicated in red. **C.** Neuro2A cells were transfected with WT HA-NFATc3 or NFATc3-EallA for 48 h. Then, cells were incubated with 20 μg/ml cycloheximide (CHX) for increasing times before harvesting. Proteins were analyzed by western blot using antibodies against HA tag and tubulin. **D.** The intensity of the bands on the western blots of different experiments performed as in C was quantified. For each experiment, the amount of NFATc3 was normalized by the level of tubulin in each condition and plotted against CHX incubation time. Data are the mean ± SEM of three independent experiments. ***p<0.0001, **p<0.005 significantly different from WT NFATc3 at the same incubation time (two-way ANOVA followed by Sidak’s multiple comparisons test).

### Trim39 acts as a SUMO-targeted E3 ubiquitin-ligase for NFATc3

To better understand the mechanisms underlying the regulation of NFATc3 by SUMO, we examined whether mutation of its three consensus SUMOylation sites could affect its ubiquitination by Trim39. Indeed, the ubiquitination level of the NFATc3-EallA mutant was decreased compared to WT NFATc3 when co-expressed with Trim39 in Neuro2A cells (Fig. 7A). To determine whether this could be due to a reduced interaction between NFATc3 and Trim39, co-immunoprecipitation experiments were performed. The amount of Trim39 co-precipitated with NFATc3-EallA was decreased compared to the amount of Trim39 co-precipitated with WT NFATc3 (Fig. 7B left panel). Consistently, the amount of NAFTc3 co-precipitated with Trim39 was decreased when its three SUMOylation sites were mutated (Fig. 7B right panel). Therefore, these data suggest that Trim39 binds and ubiquitinates preferentially SUMOylated forms of NFATc3.

**Figure 7.**
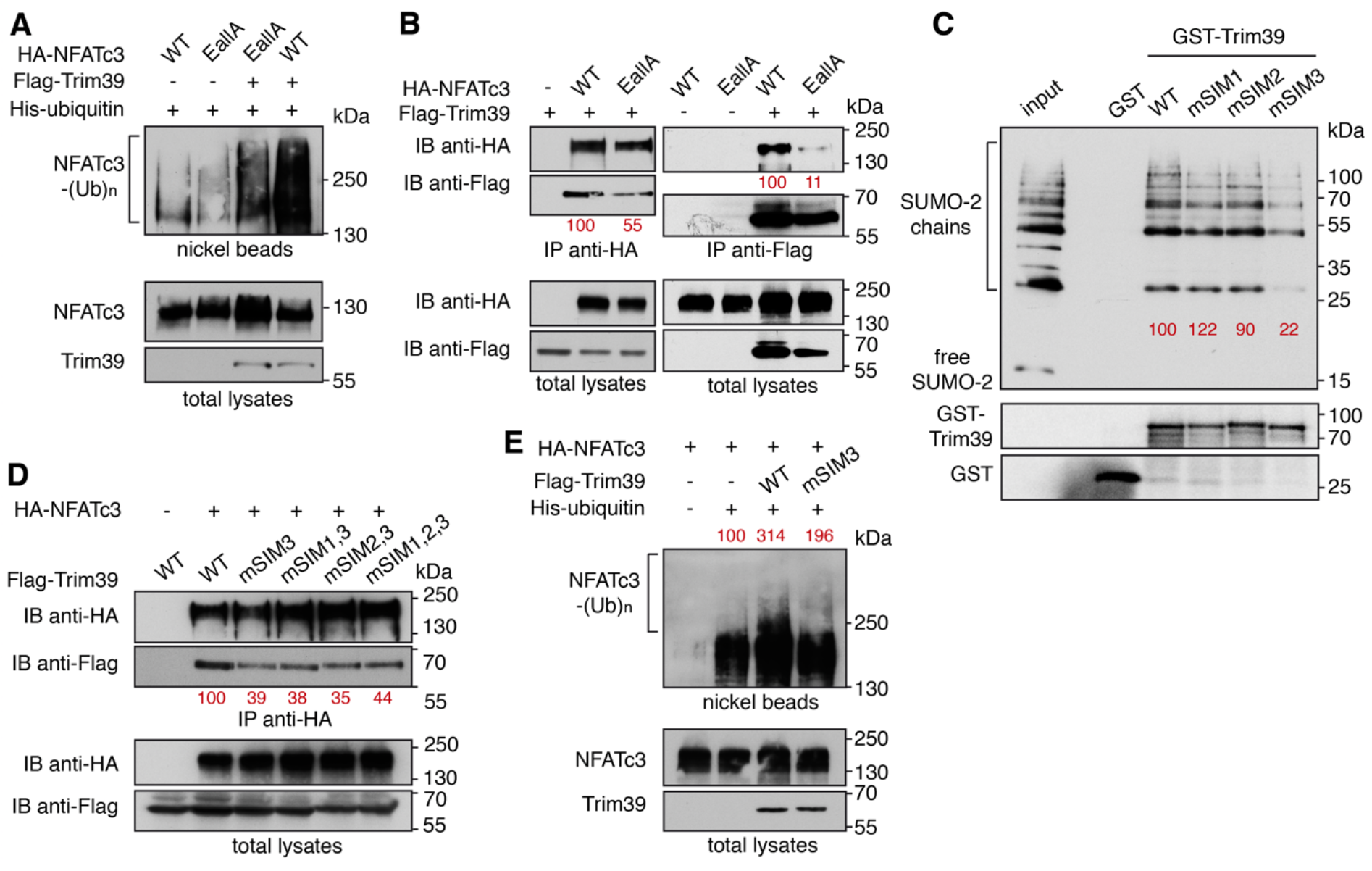
Trim39 is a SUMO-targeted E3 ubiquitin-ligase for NFATc3. **A.** Neuro2A cells were transfected with His-tagged ubiquitin together with WT HA-NFATc3 or HA-NFATc3 EallA, in the presence or the absence of Flag-Trim39, for 24 h. Then, cells were incubated with 20 μM MG-132 for 6 h before harvesting. The ubiquitinated proteins were purified using nickel beads and analyzed by western blotting using anti-HA antibody to detect ubiquitin-conjugated HA-NFATc3. In a separate SDS-PAGE, samples of the input lysates used for the purification were analyzed with antibodies against HA and Flag. **B.** Neuro2A cells were transfected with Flag-Trim39 together with WT HA-NFATc3, HA-NFATc3-EallA or empty plasmid for 24 h. Cells were then treated with 10 μM MG-132 for 8 h. The cells were subsequently harvested and lysates were subjected to immunoprecipitation using anti-HA antibody (left panel) or anti-Flag beads (right panel). Immunoprecipitates and total lysates were analyzed by western blot using anti-HA and anti-Flag antibodies. The intensity of the bands containing Flag-Trim39 co-immunoprecipitated with HA-NFATc3 was normalized by the intensity of the bands of immunoprecipitated HA-NFATc3. The intensity of the bands containing HA-NFATc3 co-immunoprecipitated with Flag-Trim39 was normalized by the intensity of the bands of immunoprecipitated Flag-Trim39. Relative values are indicated in red. **C.** Recombinant GST, GST-Trim39 and its different SIM mutants were purified using glutathione beads and subsequently incubated with purified recombinant SUMO-2 and SUMO-2 chains. Material bound to the beads was eluted and analyzed by western blot using anti-SUMO and anti-GST antibodies. A small fraction of the SUMO-2 chains was also loaded on the gel (input) for comparison. The intensity of bound SUMO-chain bands was quantified and normalized by the intensity of corresponding GST-Trim39 bands. Relative values are indicated in red. Note that SUMO bands are multiple of ≈ 15 kDa corresponding to mono-, di-, tri-, tetra-SUMO etc… **D.** Neuro2A cells were transfected with HA-NFATc3 or empty plasmid together with WT Flag-Trim39 or its SIM3 mutant for 24 h. Cells were treated as in B and lysates were subjected to immunoprecipitation using anti-HA antibody. Immunoprecipitates and total lysates were analyzed as in B. The intensity of the bands containing Flag-Trim39 co-immunoprecipitated with HA-NFATc3 was normalized by the intensity of the bands of immunoprecipitated HA-NFATc3. Relative values are indicated in red. **E.** Neuro2A cells were transfected with His-tagged ubiquitin (or empty plasmid) together with HA-NFATc3 in the presence or the absence of Flag-Trim39 or its SIM3 mutant, for 24 h. Then, cells were treated as in A. Ubiquitinated proteins and input lysates were analyzed as in A. The intensity of the ubiquitinated forms of NFATc3 was quantified and normalized by the intensity of NFATc3 in the total lysate. Relative values are indicated in red.

Proteins interacting non-covalently with SUMO generally harbor SUMO-interacting motifs. (SIMs). These motifs typically consist of three hydrophobic residues in a sequence of four amino acids, sometimes flanked by acidic or phosphorylated residues (Kerscher, 2007). Using the web-based tool GPS-SUMO (Q. Zhao et al., 2014), we identified three putative SIMs in the Trim39 sequence, which are conserved from mouse to human. We named these motifs SIM1 (39-PVII-42, located in the RING domain), SIM2 (125-VCLI-128, in the B-Box domain) and SIM3 (211-LLSRL-215, in the coiled-coil domain). These three putative SIMs exhibit the highest predictive scores with GPS-SUMO. Two of them, SIM1 and SIM2, are also predicted with a high score by the JASSA bioinformatics tool (Beauclair et al., 2015). Trim39 constructs were generated in which most residues of the three SIMs were mutated to Ala (respectively into mSIM1: 39-PAAA-42, mSIM2: 125-AAAA-128 and mSIM3: 211-AAARA-215). To confirm the ability of Trim39 to bind SUMO and to determine the impact of these mutations, we conducted GST pull-down experiments using purified recombinant proteins. Interestingly, GST-Trim39 could bind di-, tri-, tetra- and higher-order SUMO-2 chains but not free SUMO-2, whereas GST alone showed no interaction (Fig. 7C). Single mutations of SIM1 and SIM2 had no significant effect, either individually (Fig. 7C) or together (Fig. S3). In contrast, mutation of SIM3 strongly reduced the SUMO-binding ability of Trim39 (Fig. 7C), an effect which was not significantly modified by combination with single SIM1 mutation, and only slightly increased by combination with single SIM2 and double SIM1/SIM2 mutations (Fig. S3), as previously described for the SUMO-target E3 ubiquitin-ligase Arkadia/RNF11 (Erker et al., 2013). These data suggest that SIM3 plays a pivotal role in the binding of Trim39 to SUMO chains. Consistently, mutation of SIM3 reduced the ability of Trim39 to interact with NFATc3 in co-immunoprecipitation experiments (Fig. 7D). As for SUMO-2 chain binding (Fig. 7C), the concomitant mutation of SIM1, SIM2 or both, together with SIM3, did not significantly modify the binding of Trim39 to NFATc3 (Fig. 7D). Moreover, SIM3 mutation reduced the ability of Trim39 to ubiquitinate NFATc3 in Neuro2A cells (Fig. 7E).

Collectively, these data strongly suggest that Trim39 acts as a SUMO-targeted E3 ubiquitin-ligase for NFATc3 by preferentially binding the SUMOylated forms of NFATc3 through its SIM, in order to mediate their ubiquitination.

### SUMOylation and Trim39 modulate the pro-apoptotic effect of NFATc3 in neurons

In a previous study, we have shown that overexpression of NFATc3 in primary cultures of cerebellar granule neurons (CGNs) aggravates apoptosis induced by KCl deprivation (Mojsa et al., 2015). Primary CGNs represent one of the best characterized *in vitro* models of neuronal apoptosis (Contestabile, 2002).These neurons survive in the presence of serum and depolarizing concentrations of KCl (25 mM) that mimic the neuronal activity required for their survival *in vivo* (Ikonomidou et al., 1999). They undergo apoptosis following withdrawal of serum and lowering of KCl to 5 mM (K5) (D’Mello et al., 1993), which recapitulates the programmed cell death naturally occurring in the cerebellum during post-natal development (Wood et al., 1993). We used this model to examine whether mutation of the SUMOylation sites of NFATc3, which increases its stability (Fig. 6D) by reducing its interaction with Trim39 (Fig. 7B) and its ubiquitination (Fig. 7A), could have an impact on its pro-apoptotic effect in CGNs. As shown previously (Mojsa et al., 2015), we confirmed that KCl deprivation-induced apoptosis is significantly increased in CGNs transfected with WT GFP-NFATc3 compared to GFP, as shown by the increased number of apoptotic/condensed nuclei (Fig. 8A,B). Interestingly, neuronal apoptosis was further increased in neurons overexpressing GFP-NFATc3-EallA compared to WT GFP-NFATc3 (Fig. 8A,B). Consistently, efficient silencing of Trim39 using a lentivirus expressing a specific shRNA (Fig. 8C) significantly aggravated apoptosis compared to neurons transduced with an unrelated control shRNA (Fig. 8D,E). Our data therefore strongly suggest that SUMO and Trim39 negatively regulate the pro-apoptotic function of NFATc3, most likely by reducing its stability and thereby its activity as a transcription factor.

**Figure 8.**
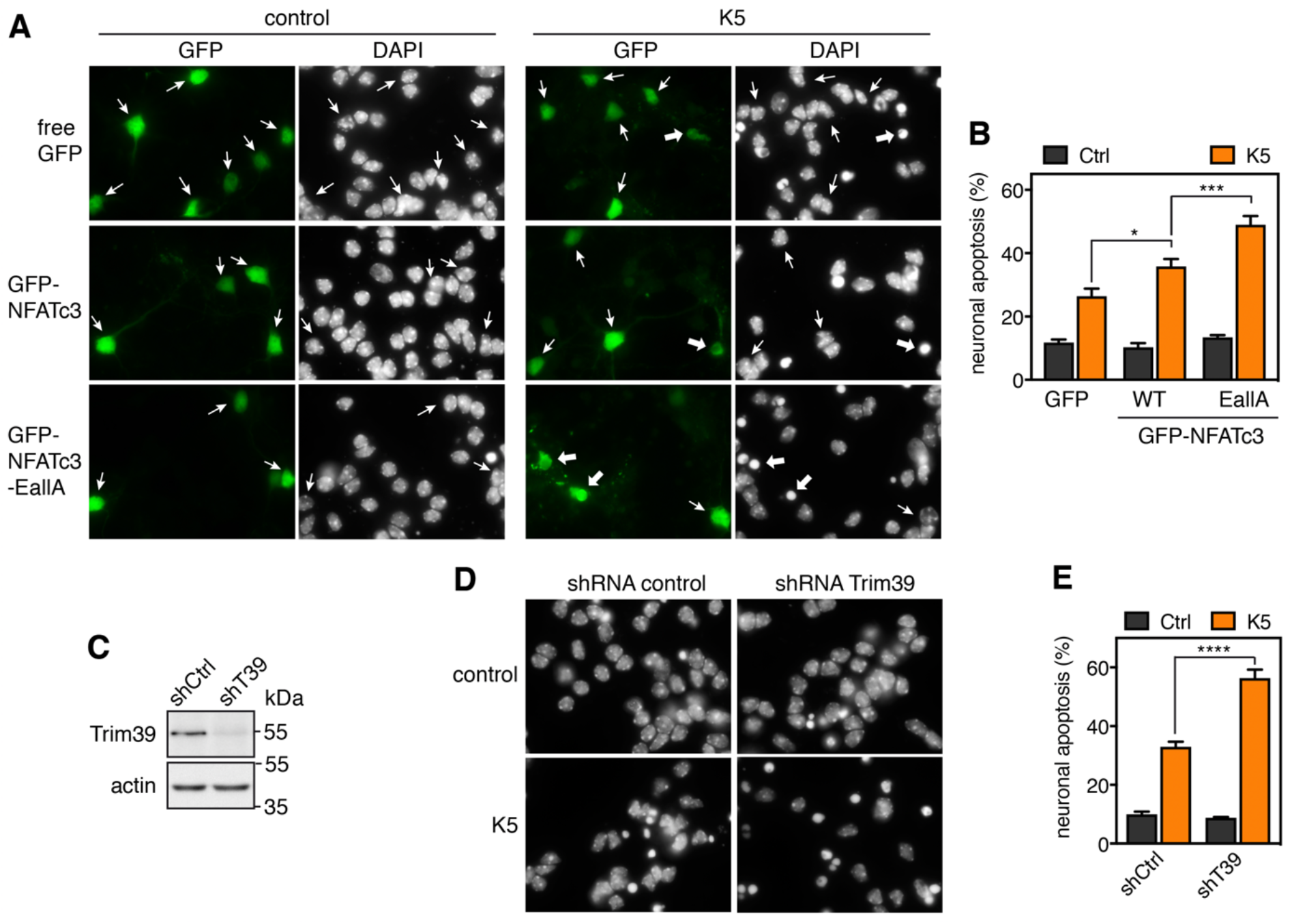
SUMOylation and Trim39 attenuate NFATc3 pro-apoptotic effect in neurons. **A.** CGN primary cultures were transfected after 5 days *in vitro* (DIV 5) with GFP (as a negative control), WT GFP-NFATc3 or GFP-NFATc3-EallA for 16 h. Then, neurons were switched to serum-free medium containing 5 mM KCl (K5) for 7 h or were left untreated (control). Following fixation, nuclei were visualized by DAPI staining and proteins fused to GFP were detected by fluorescent microscopy. Arrows indicate GFP-positive neurons with thick arrows for neurons undergoing apoptosis and thin arrows for healthy neurons. **B.** The percentage of transfected, GFP-positive neurons undergoing apoptosis was assessed by examining cell morphology and nuclear condensation. Data are the means ± S.E.M. of four independent experiments performed as in A. * *P*<0.05; *** * *P*<0.001 significantly different from the corresponding value obtained in neurons transfected with GFP (two-way ANOVA followed by Sidak’s multiple comparisons test). **C.** CGNs were transduced with lentiviral particles expressing a non-targeting control (directed against Luciferase) or an shRNA specifically targeting Trim39 one day after plating. At DIV 6, total cell extracts from KCl-deprived neurons were analyzed by western blot using anti-Trim39 antibody (Origene). **D.** CGN were transduced and treated as in C. At DIV 6 they were incubated for 8 h in K5 medium, fixed and stained with DAPI. **E.** The percentage of apoptotic neurons was estimated by examining nuclear condensation. Data are the means ± S.E.M. of four independent experiments performed as in D. **** *P*<0.0001 significantly different from neurons transduced with the control shRNA (two-way ANOVA followed by Sidak’s multiple comparisons test).

## Discussion

In contrast to calcium/calcineurin-mediated NFAT nuclear translocation, the regulation of NFAT protein stability by the ubiquitin-proteasome system has been poorly studied. Independent studies have suggested that certain E3 ubiquitin-ligases may be responsible for ubiquitination and proteasomal degradation of different NFAT members: HDM2 for NFATc2 in breast and pancreatic cancers (Singh et al., 2011; Yoeli-Lerner et al., 2005); Cbl-b, c-Cbl, VHL or KBTBD11/Cullin3 for NFATc1 during osteoclastogenesis (J. H. Kim et al., 2010; X. Li et al., 2015; Narahara et al., 2019; Youn et al., 2012); CHIP for NFATc3 in LPS-induced cardiomyopathies (Chao et al., 2019). However, no formal demonstration has been made to establish that these proteins are genuine NFAT E3 ubiquitin-ligases, with the exception of HDM2 for NFATc2 (Yoeli-Lerner et al., 2005). In the present study, we provide the first formal identification of an NFATc3 E3 ubiquitin-ligase. Indeed, we show several lines of evidence demonstrating that Trim39 is indeed an E3 ubiquitin-ligase for NFATc3. First, we found a physical interaction between endogenous or overexpressed Trim39 and NFATc3 proteins. Second, Trim39 ubiquitinated NFATc3 *in vitro*. Third, overexpression of WT Trim39, but not of its inactive RING mutant, increased the ubiquitination level of NFATc3 in cells. In contrast, silencing of Trim39 decreased NFATc3 ubiquitination. Finally, Trim39 overexpression decreased the protein level of NFATc3 whereas the silencing of endogenous Trim39 increased it, suggesting that Trim39-mediated ubiquitination of NFATc3 targets it for proteasomal degradation. As a physiological consequence, overexpressed Trim39 resulted in reduced transcriptional activity of NFATc3 without affecting its nuclear translocation. Conversely, silencing of endogenous Trim39 increased both the expression of a NFATc3 target gene and its pro-apoptotic effect in neurons. Taken together, these data strongly suggest that Trim39 modulates neuronal apoptosis by acting as a physiological E3 ubiquitin-ligase for NFATc3. This does not exclude the possibility that other NFATc3 E3 ubiquitin-ligases exist, notably in other cell types, such as CHIP in cardiomyocytes (Chao et al., 2019). Nevertheless, NFATc3 ubiquitination is deeply decreased following Trim39 knock-down in Neuro2A cells, suggesting that Trim39 is the major E3 ubiquitin-ligase for NFATc3 in these cells.

Our present data show that Trim17 inhibits the ubiquitination of NFATc3 mediated by Trim39. Indeed, the increase in the ubiquitination level of NFATc3 due to overexpression of Trim39 was abolished by the co-transfection of Trim17. Both Trim39 and Trim17 belong to the family of TRIM proteins which forms one of the largest classes of RING-containing E3 ubiquitin-ligases (Meroni & Diez-Roux, 2005), comprising 82 members in humans (Qiu et al., 2020). TRIM proteins are characterized by their N-terminal tripartite motif that consists of a RING domain, one or two B-box domains and a coiled-coil domain, invariably ordered in this sequence from N- to C-termini (Esposito et al., 2017; Reymond et al., 2001). In addition to this common motif, TRIM proteins generally exhibit a C-terminal domain that varies from one member to another and categorizes them into different subtypes (Short & Cox, 2006). This C-terminal domain, which is a PRY-SPRY domain for both Trim17 and Trim39, generally mediates target recognition and specificity (Esposito et al., 2017; Y. Li et al., 2014). While the RING domain confers an E3 ubiquitin-ligase activity by binding ubiquitin-loaded E2 ubiquitin-conjugating enzymes, the B-box and especially the coiled-coil domain are involved in the formation of homo- or hetero-dimers or multimers (Koliopoulos et al., 2016; Y. Li et al., 2014; Napolitano & Meroni, 2012; Sanchez et al., 2014). As homo-oligomerization seems to be necessary for the E3 ubiquitin-ligase activity of TRIM proteins (Koliopoulos et al., 2016; Streich et al., 2013; Yudina et al., 2015), Trim17 may inhibit Trim39-mediated ubiquitination of NFATc3 by forming inactive hetero-oligomers with Trim39. Indeed, we show here that Trim17 and Trim39 physically interact with each other. In a similar way, we have previously shown that TRIM17 inhibits the activity of two other TRIM proteins to which it is able to bind: TRIM41 (Iréna Lassot et al., 2018) and TRIM28 (Lionnard et al., 2019). It is interesting to note that TRIM39 and TRIM41 are very close from a phylogenetic point of view (Qiu et al., 2020; Sardiello et al., 2008), suggesting that common mechanisms could be involved in their inhibition by Trim17.

The inhibitory effect of Trim17 might result from two mechanisms that are not mutually exclusive. First, the formation of hetero-dimers or hetero-multimers with Trim17 may inhibit the intrinsic E3 ubiquitin-ligase activity of its TRIM partner, possibly by preventing the binding of the E2 enzyme. Indeed, we show in the present study that TRIM17 prevents the auto-ubiquitination of TRIM39 *in vitro*, similarly to what we have previously shown for the inhibition of TRIM41 by TRIM17 (Iréna Lassot et al., 2018). Second, Trim17 may prevent the binding of the substrate to its TRIM partner. Indeed, we show here that Trim17 reduces the interaction between Trim39 and NFATc3, as determined by both co-IP of overexpressed proteins and proximity ligation of endogenous proteins (PLA). These results are similar to the effect of TRIM17 on the interaction between TRIM41 or TRIM28 and their respective substrates (Iréna Lassot et al., 2018; Lionnard et al., 2019). In the three cases, Trim17 is able to bind both the substrate and the TRIM partner, suggesting that Trim17 could reduce their interaction either by directly binding the substrate in a competitive manner, or because the formation of a hetero-dimer hinders the accessibility of the TRIM partner to the substrate. Further experiments are required to determine the structural determinants of the inhibitory effect of Trim17 on other TRIM proteins. Nevertheless, it is unlikely that Trim17 inhibits Trim39-mediated ubiquitination of NFATc3 by associating with a deubiquitinating enzyme (DUB), as shown for other TRIM proteins (Hao et al., 2015; Nicklas et al., 2019). Indeed, TRIM17 is able to inhibit the *in vitro* ubiquitination of NFATc3 mediated by TRIM39, in a completely acellular medium in the absence of any DUB. It is also clear that Trim17 does not inhibit Trim39 by inducing its ubiquitination and subsequent degradation. Indeed, Trim17 rather decreases the ubiquitination level of Trim39 both *in vitro* and in cells. Moreover, it is interesting to note that, TRIM39 reciprocally decreases the *in vitro* auto-ubiquitination of TRIM17, further suggesting that Trim17 and Trim39 form inactive hetero-dimers or hetero-multimers, in which the E3 ubiquitin-ligase activity of the two partners is inhibited. Formation of inactive hetero-oligomers might be favored by the fact that Trim17 and Trim39 belong to the same phylogenetic group (Qiu et al., 2020; Sardiello et al., 2008), as hetero-interactions are more often observed among closely related TRIM family members (Napolitano & Meroni, 2012).

SUMOylation of proteins changes their non-covalent interactions in various ways, leading to alterations in their subcellular localization and function. As such, SUMO modification plays crucial roles in a wide range of cellular processes (Henley et al., 2018; Yau et al., 2020; X. Zhao, 2018). SUMOylation has also emerged as an important regulator of protein stability (Liebelt & Vertegaal, 2016). Consistently, our data clearly indicate that SUMOylation of NFATc3 promotes its ubiquitination and subsequent degradation. Indeed, mutation of the three SUMOylation consensus sites of NFATc3 decreased its ubiquitination level, increased its half-life and aggravated its pro-apoptotic effect in neurons. As the SUMOylation sites were modified in order to specifically prevent SUMOylation without affecting a putative ubiquitination of the acceptor Lys residues, our results unambiguously designate SUMOylation as the post-translational modification implicated in these effects. This mechanism could be conserved in other members of the NFAT family. Indeed, effective SUMOylation of NFAT proteins has been described, notably for NFATc1 and NFATc2 (Nayak et al., 2009; Terui et al., 2004). Although the functional consequences of NFAT SUMOylation that have been reported so far are rather related to their nuclear translocation and transactivation activity (E. T. Kim et al., 2019; Nayak et al., 2009; Terui et al., 2004; Vihma & Timmusk, 2017), it might also influence the stability of these proteins. Indeed, Singh et al. reported that the double mutation of Lys684 and Lys897 in murine NFATc2 prevents its ubiquitination and degradation induced by zoledronic acid (Singh et al., 2011). One possible conclusion is that NFATc2 is normally ubiquitinated on these Lys residues (Singh et al., 2011). However, as these two Lys residues have been shown to be SUMOylated (Terui et al., 2004), an alternative plausible explanation is that SUMOylation of NFATc2 on these consensus sites might be necessary for the recognition by its E3 ubiquitin-ligase. In line with this hypothesis, the protein level of different human NFATc1 and NFATc2 isoforms have been reported to be increased by the double K/R mutation (lysine to arginine substitution) of their C-terminal SUMOylation sites (Vihma & Timmusk, 2017). However, further investigation is required to demonstrate that SUMO indeed plays a role in these effects.

A few SUMO-targeted E3 ubiquitin-ligases (STUbLs) have been described (Geoffroy & Hay, 2009; Prudden et al., 2007; Sriramachandran & Dohmen, 2014). These proteins generally combine two features: a RING domain, that confers them an E3 ubiquitin-ligase activity, and SUMO interacting motifs (SIMs) that mediate their preference for SUMOylated substrates (Geoffroy & Hay, 2009; Prudden et al., 2007; Sriramachandran & Dohmen, 2014). Inhibition of the proteasome leads to an important accumulation of high molecular weight SUMO-modified proteins in yeast and human cells (Bailey & O’Hare, 2005; Uzunova et al., 2007), suggesting that ubiquitination and degradation of SUMOylated proteins mediated by STUbLs play an important role in proteostasis. However, only two STUbLs have been identified so far in mammals: RNF4 and Arkadia/RNF111 (Branigan et al., 2019; Sriramachandran & Dohmen, 2014), which may not be sufficient to account for the regulation of all SUMOylated proteins. Therefore, additional STUbLs probably remain to be discovered. Here we provide a series of arguments indicating that Trim39 acts as a STUbL for NFATc3. First, Trim39 is able to bind SUMO chains but not free SUMO *in vitro*. Second, we identified a SIM in the sequence of Trim39. Indeed, mutation of this motif strongly decreases its capacity to bind SUMO-2 chains *in vitro*. Third, the binding and the ubiquitination of NFATc3 mediated by Trim39 is reduced by mutation of this SIM in Trim39. Finally, Trim39 binds and ubiquitinates preferentially the SUMOylated forms of NFATc3.

Most of the STUbLs studied so far bear multiple SIMs, which mediate cooperative binding to multiple SUMO units, thereby providing a preference for substrates with SUMO chains (Sriramachandran & Dohmen, 2014). Among the three SIMs predicted in the Trim39 sequence with a high score, only one is really involved in the binding of Trim39 to purified SUMO chains and SUMOylated NFATc3 in cells. SIMs are characterized by a loose consensus sequence and some non-canonical SIMs have been described (Kerscher, 2007; Sriramachandran & Dohmen, 2014). Therefore, it is possible that another SIM in Trim39, that was not identified or was not credited with a high predictive score by GPS-SUMO or JASSA, may participate in the binding of Trim39 to SUMO. It is also possible that one SIM is sufficient to fulfill this function, as has been reported for other STUbLs (Erker et al., 2013; Parker & Ulrich, 2012). For example, among the three SIMs identified in Arkadia/RNF111, only one has been shown to be crucial for its interaction with SUMO-2 chains, exactly like SIM3 in Trim39 (Erker et al., 2013). Moreover, the activity of TRIM proteins generally involves homo-dimerization or homo-multimerization (Koliopoulos et al., 2016; Streich et al., 2013; Yudina et al., 2015). It is therefore possible that in its multimerized form, Trim39 harbors several SIMs in close proximity. Indeed, SIM3 is located in the coiled-coil domain of Trim39 that is expected to form antiparallel dimers or higher-order multimers, as reported in other TRIM proteins (Koliopoulos et al., 2016; Y. Li et al., 2014; Napolitano & Meroni, 2012; Sanchez et al., 2014). Therefore, the unique active SIM of one molecule of Trim39 may cooperate with the SIM of other Trim39 molecules, in multimers, for binding SUMO chains. Alternatively, another binding site, such as the RING domain, may cooperate with the SUMO-SIM interaction to bind the substrate, as has been reported for the *Drosophila* STUbL Degringolade (Abed et al., 2011). Trim39 may also bind NFATc3 by a dual mechanism, involving both SUMO-dependent and SUMO-independent recognition, as shown for viral STUbLs (Boutell et al., 2011; Wang et al., 2011). Indeed, we found that the NFATc3/Trim39 interaction is decreased but not completely abrogated, in co-immunoprecipitation experiments, by mutation of either the SUMOylation sites in NFATc3 or SIM3 in Trim39. Further studies are required to fully characterize the mechanisms mediating the SUMO-dependent interaction of NFATc3 with Trim39.

TRIM proteins play significant roles in a wide variety of cellular processes including cell cycle, differentiation, apoptosis, autophagy, transcription, DNA repair, innate immunity and anti-viral defense (Hatakeyama, 2017; Venuto & Merla, 2019). As a consequence, mutations of *TRIM* genes have been implicated in several human diseases such as cancer, auto-immune diseases, rare genetic disorders, infectious diseases and neurodegenerative diseases (Hatakeyama, 2017; Meroni, 2020; Watanabe & Hatakeyama, 2017). In particular, Trim39 has been shown to regulate cell cycle progression by directly mediating the ubiquitination of p53 (Zhang, Huang, et al., 2012) and by indirectly modulating the protein level of p21 (Zhang, Mei, et al., 2012). Trim39 has also been implicated in the negative regulation of NFκB signaling (Suzuki et al., 2016). In the present study, we show that silencing of Trim39 enhances apoptosis triggered by survival-factor deprivation in primary cultures of neurons. As Trim39 induces NFATc3 degradation and as NFATc3 aggravates neuronal apoptosis, these data suggest that silencing of Trim39 favors apoptosis of neurons by stabilizing NFATc3. However, it does not exclude additional effects of Trim39 on apoptosis regulation. For example, silencing of Trim39 has been reported to increase nutlin3a-induced apoptosis in several p53-positive cancer cell lines, presumably by stabilizing the pro-apoptotic factor p53 (Zhang, Huang, et al., 2012). Trim39 knock-down has also been reported to aggravate apoptosis following a genotoxic stress in HCT116 cells (Zhang, Mei, et al., 2012). Conversely, in p53 null cell lines, silencing of Trim39 reduces DNA damage-induced apoptosis (S. S. Lee et al., 2009; Zhang, Huang, et al., 2012). The latter effect is likely due to the ability of Trim39 to directly inhibit APC/C^Cdh1^-mediated degradation of the pro-apoptotic protein MOAP-1 (Huang et al., 2012; S. S. Lee et al., 2009). These data, taken together with the present study, place Trim39 as a key factor in several processes regulating apoptosis, the final outcome depending on its targets or binding partners present in the cell and the nature of the cellular stress. Similarly, we and others have reported that Trim17 plays an important role in apoptosis regulation (I. Lassot et al., 2010; Lionnard et al., 2019; Magiera et al., 2013; Mojsa et al., 2015; Song et al., 2017). Notably, we have shown that Trim17 is both sufficient and necessary to trigger apoptosis in primary neurons (I. Lassot et al., 2010), at least in part by mediating the ubiquitination and subsequent degradation of the anti-apoptotic protein Mcl-1 (Magiera et al., 2013). Trim17 may also modulate neuronal apoptosis by acting on NFATc3 through antagonistic mechanisms. On one hand, we have previously shown that Trim17 can prevent the nuclear translocation and transcriptional activity of NFATc3 (Mojsa et al., 2015) and should therefore inhibit its pro-apoptotic effect. On the other hand, we show here that Trim17 can inhibit Trim39-mediated ubiquitination and degradation of NFATc3 and should therefore aggravate its pro-apoptotic effect by increasing its protein level. Moreover, we have identified *Trim17* as a target gene of NFATc3 (Mojsa et al., 2015). The effects of Trim17 on the protein level and activity of NFATc3 should therefore influence its own expression, creating both a negative and a positive feedback loop and eventually resulting in fine tuning of neuronal apoptosis. Further investigation will determine whether these mechanisms can be manipulated for therapeutic purposes to prevent neuronal loss in neurodegenerative diseases.

## Materials and Methods

### Materials

Culture media were from Thermo Fisher Scientific. Fetal calf serum, other culture reagents, protease inhibitor cocktail, DAPI, PMA, A23187, cycloheximide, N-ethylmaleimide (NEM), MG-132, puromycin, anti-Flag M2 affinity gel beads (#A2220) and other chemicals were from Sigma-Aldrich. Protein G-agarose and protein A-agarose beads were from Roche. GFP-Trap®-A beads were from Chromotek (Planneg-Martinsried, Germany). Rat monoclonal anti-HA antibody (clone 3F10; #11867432001), mouse monoclonal anti-Flag antibody (clone M2, #F3165), mouse monoclonal anti-tubulin antibody (clone DM1A, #T6199), rabbit anti-TRIM17 antibody (#AV34547) and rabbit IgG (#I5006) were from Sigma-Aldrich. Rabbit anti-GFP antibody (#TP401) was from Torrey Pines Biolabs Inc. (Houston, TX USA). Mouse monoclonal antibody against actin (clone C4) was from Millipore (#MAB1501). Rabbit polyclonal antibody against NFATc3 was from Proteintech (18222-1-AP). Mouse monoclonal antibody against Trim39 was from Origene (#TA505761). Rabbit polyclonal antibody against Trim39 was from Proteintech (#12757-1-AP). Monoclonal mouse antibody against SUMO-2 (clone #8A2) was purified from hybridomas obtained from the Developmental Studies Hybridoma Bank. Fluorescent and horseradish peroxidase-conjugated goat anti-rabbit, anti-rat and anti-mouse secondary antibodies were from Thermo Fisher Scientific and Jackson ImmunoResearch Laboratories Inc. (West Grove, PA, USA), respectively.

### Cell culture and transient transfection

Lenti-X 293 T (Clontech), Neuro2A (mouse neuroblastoma) and Baby Hamster Kidney (BHK) cell lines were grown in Dulbecco’s modified Eagle’s medium containing 4.5 g/l glucose supplemented with 10% fetal bovine serum and penicillin-streptomycin 100 IU/ml-100 μg/ml. Cells were transfected with plasmids using GenJet™ *in vitro* transfection reagent (Ver. II) pre-optimized and conditioned for transfecting Neuro2A and BHK-21 cells respectively (SignaGen laboratories, Ijamsville, MD) according to the manufacturer’s instructions. Neuro2A cells were transfected with siRNAs using Lipofectamine RNAiMAX transfection reagent (Thermo Fisher Scientific) following the manufacturer’s instruction. For one 35 mm dish, 2.5 μl of transfection reagent was used with 25 pmoles of siRNA. The sequences of the siRNAs used were as follows: siTrim39#1 5’ CCAAGCGGGTAGGCATATT 3’; siTrim39#2 5’ GCGTCAAGTTTGTGGAGACAA3’; siRNA ctrl (targeting Luciferase gene): 5’ CGTACGCGGAATACTTCGA 3’.

Primary cultures of CGNs were prepared from 7-day-old murine pups (C57Bl/6 J mice) as described previously (I. Lassot et al., 2010). Briefly, freshly dissected cerebella were dissociated by trypsinization and mechanical disruption, and plated in Basal Medium Eagle (BME) medium supplemented with 10% fetal bovine serum, 2 mM L-Gln, 10 mM HEPES, penicillin-streptomycin 100 IU/ml-100 μg/ml and 20 mM KCl. Primary CGNs, grown on glass coverslips in 24-well plates, were transfected at DIV 5 with 2 μg of plasmids using a calcium phosphate protocol optimized for neuronal cultures as previously described (I. Lassot et al., 2010).

### Silencing of Trim39 using shRNA-expressing lentiviruses

The HIV-derived lentiviral vectors pLKO.1 containing control shRNAs respectively against eGFP and Luciferase (SHC005, SHC007) and the shRNAs TRCN0000037281 (shRNA Trim39#1), TRCN0000037282 (shRNA Trim39#2) and TRCN0000438509 (shRNA Trim39#3) were from Sigma-Aldrich. Lentiviral particles were produced as previously described (Iréna Lassot et al., 2018). Neuro2A cells were transduced one day after plating. The lentiviral preparations were added directly to the culture medium for 8 h (approximately 500 ng p24 per million neurons, approximately 100 ng p24 per million Neuro2A cells). Cells were then replaced in fresh medium. Culture was continued until 6 days in vitro for CGNs. Neuro2A cells were maintained in culture for 24 h after transduction and then selected using 2 μg/ml puromycin for an additional 48 h.

### Expression vectors and site directed mutagenesis

The following plasmids were described previously: pCI-GFP, pCS2-3×HA-NFATc3, pCS2-3×HA-NFATc3-KallR, pCS2-GFP-NFATc3 and pCI-Trim17-GFP (Mojsa et al., 2015). All the primers used to generate the constructs described below are listed in supplementary Table1. The sequences of all the constructs were confirmed by automatic sequencing. Single point mutations in the SUMOylation-consensus sites of NFATc3 (E437A, E706A and E1015A) or their double and triple combinations (E437A/E706A, E706A/E1015A and E437A/E706A/E1015A=EallA) were obtained by site-directed mutagenesis of pCS2-3×HA-NFATc3 using the QuickChange® II XL kit (Agilent Technologies) using the indicated primers. To increase the expression of NFATc3, HA-NFATc3 and HA-NFATc3-EallA from respective pCS2-3×HA expression vectors were first sub-cloned between *XhoI* and *NheI* sites of the pCDNA3.1 plasmid, and then sub-cloned between *SalI* and *NheI* sites of the pCI plasmid, to generate pCI-3×HA-NFATc3 and pCI-3×HA-NFATc3-EallA. The plasmid pCS2-GFP-NFATc3-EallA was obtained by removing the WT NFATc3 cDNA from the plasmid pCS2-GFP-NFATc3 and by replacing it with NFATc3-EallA between *EcoRI* and *XhoI* sites of the plasmid. The cDNA of mouse Trim39 (GenBank: NM_024468) was amplified, from primary CGN cDNAs, by using PCR with the indicated primers. Amplicons were then cloned into pCI-3×Flag plasmid between *EcoRI* and *XbaI* sites to obtain mouse pCI-3×Flag-Trim39. pGEX-4T1-Trim39-ΔRING mutant was generated by PCR amplification of Trim39 coding region using pCI-3×Flag-Trim39 as template and the indicated primers. Then, the amplicons were cloned between *EcoRI* and *XhoI* sites of the plasmid pGEX4T1. To obtain pCI-3×Flag-Trim39-ΔRING, the insert was released from the plasmid pGEX4T1-Trim39-ΔRING and sub-cloned between *EcoRI* and *NotI* sites of the plasmid pCI-3×Flag. The following Trim39 SIM mutants were obtained by site-directed mutagenesis using pCI-3×Flag-Trim39 as a template and the indicated primers: single mutants mSIM1 (PVII→PAAA), mSIM2 (VCLI→ACAA) and mSIM3 (LLSRL→AAARA); double mutants mSIM12, mSIM13 and mSIM23; and triple mutant mSIM123. The expression vector pmCherry-C1 was purchased from Takara Bio Inc. (#632524). The plasmid mouse pCI-Trim39-mCherry was obtained by recombinant PCR. The first PCR was performed with the indicated primers using pCI-3×Flag-Trim39 as a template. The second PCR was performed with the indicated primers using pmCherry-C1 as a template. The amplicons from both PCRs were purified, mixed and used as template for the recombinant PCR (third PCR) with the indicated primers. The resulting amplicon was cloned between *EcoRI* and *XbaI* sites of the empty pCI plasmid to obtain pCI-Trim39-mCherry.

In order to produce recombinant N-terminal GST-tagged Trim39 protein in *Escherichia coli*, the pGEX-4T1-Trim39 construct was produced by PCR amplification of the Trim39 coding region using pCI-3×Flag-Trim39 as a template and the indicated primers. The PCR products were cloned between the *EcoRI* and *XhoI* sites of the pGEX-4T1 expression vector (GE-Healthcare). The mutant GST-Trim39-C49S/C52S was generated by site-directed mutagenesis using GST-Trim39 as a template and the indicated primers. SIM1, SIM2 and SIM3 GST-Trim39 mutants were generated by site-directed mutagenesis using GST-Trim39 as a template and the same primers as for pCI-3×Flag-Trim39 SIM mutants. The cDNA of human TRIM39 (GenBank: NM_172016.2), N-Terminally fused to a histidine tag in the plasmid pET-15, and the cDNA of human TRIM17 (GenBank: NM_016102), C-terminally fused to GFP in the pEGFP plasmid were obtained from the ORFeome library of the Montpellier Genomic Collections facility. Human TRIM17 was first amplified by PCR using p-TRIM17-EGFP as a template and indicated primers, and the amplicons were cloned between the *EcoRI* and *SalI* sites of the pGEX-4T1 expression vector. In order to produce recombinant N-terminally MBP-tagged TRIM17 protein in *Escherichia coli*, the pMAL-c2-TRIM17 plasmid was generated by subcloning. The insert was released from the plasmid pGEX-4T1-TRIM17 and sub-cloned between the *EcoRI* and *SalI* sites of the plasmid pMAL-c2 to obtain pMAL-c2-TRIM17.

### Protein extraction and western blot analysis

Cells were harvested in lysis buffer A (50 mM Tris-HCl [pH 7.5], 150 mM NaCl, 10 mM NaF, 5 mM sodium pyrophosphate, 25 mM β-glycerophosphate, 5 mM EDTA, 10 mM NEM, 20 μM MG-132, and protease inhibitor cocktail) supplemented with 1% NP-40 and homogenized by thorough vortexing. Cell debris were removed by centrifugation at 1000 × g for 5 min at 4°C and the protein concentration of the resulting supernatant was estimated using the BCA protein assay kit (Thermo Fisher Scientific) with bovine serum albumin as the standard. Total lysates were diluted in 3 × Laemmli sample buffer and incubated at 95°C for 5 min. Proteins were separated by 8% to 12% SDS–PAGE and transferred to Immobilon-P PVDF membrane (Millipore). Blocking and probing with antibodies were performed as previously described (Iréna Lassot et al., 2018). Visualization of immunoreactive proteins was performed using horseradish peroxidase-linked secondary antibodies and Covalight enhanced chemiluminescent substrate (Covalab, Bron, France) or Immobilon® Western (Millipore). Membranes were revealed using films or Amersham Imager 680 (GE Healthcare). When necessary, membranes were stripped using Restore™ PLUS Western Blot Stripping Buffer (Thermo Fisher Scientific) and re-probed with additional antibodies.

### Co-immunoprecipitation

Following transfection with the indicated plasmids for 24 h, Neuro2A or BHK cells were incubated for 5 h with 10-20 μM MG-132. They were then homogenized in lysis buffer A, supplemented with 1% NP-40 for immunoprecipitation with anti-HA, 0.5% NP-40 for immunoprecipitation with GFP-Trap-A and 1% Triton X-100 for immunoprecipitation with anti-Flag. For anti-HA and anti-Flag immunoprecipitation, cell lysates (500 μl) were centrifuged at 300 × g for 5 min at 4°C. The resulting supernatants were pre-cleared by rotation for 1-3 h at 4°C with 15 μl protein G-agarose beads and then rotated overnight at 4°C with 25 μl protein G agarose beads together with 7 μl anti-HA antibody or with 30 μl anti-Flag M2 affinity gel beads. The beads were recovered by centrifugation and washed four times with 1 ml of lysis buffer A without detergent and containing 0,3 M NaCl for anti-HA or 0,5M NaCl for anti-Flag (instead of 150 mM NaCl). For GFP-Trap precipitation, cell lysates (200 μl) were diluted with 300 μl dilution buffer (10 mM Tris-HCl [pH 7.5], 150 mM NaCl, 10 mM NaF, 5 mM sodium pyrophosphate, 25 mM β-glycerophosphate, 0.5 mM EDTA, 20 μM MG-132, 10 mM NEM and protease inhibitor cocktail) and cell debris were removed by centrifugation. Resulting supernatants were rotated for 2 h at 4°C with 10-25 μl GFP-Trap®-A beads to immunoprecipitate proteins fused to GFP. Beads were recovered by centrifugation and washed four times with dilution buffer. Material bound to the protein G agarose, anti-Flag M2 affinity gel beads or GFP-Trap beads was eluted by the addition of 3 × Laemmli sample buffer and incubation at 95°C for 5 min. Precipitated proteins were analyzed by western blot.

### In situ proximity ligation assay

Neuro2A cells seeded onto glass coverslips, were left untreated or transfected with pCI-Flag-Trim39, pCI-GFP or pCI-Trim17-GFP for 24 h, and then treated with 10 μM MG-132 for 4 h. Cell were fixed with 4% paraformaldehyde for 20 min at room temperature, washed with 0.1 M Gly (pH = 7.11), permeabilized with 0.2 % Triton X-100 in PBS for 10 min and washed with PBS. The interaction between endogenous Trim39 and endogenous NFATc3, in the presence or absence of GFP or Trim17-GFP, was detected using the Duolink® In Situ kit (Olink® Bioscience, Uppsala, Sweden), according to the manufacturer’s instructions, as described previously (Mojsa et al., 2015). Briefly, cells were successively incubated with blocking solution for 60 min at 37°C, with primary antibodies against Trim39 (Origene, 1:400) and NFATc3 (Proteintech, 1:200) overnight at 4°C and with secondary antibodies conjugated with oligonucleotides (PLA probe MINUS and PLA probe PLUS) for 1 h at 37°C. The cells were then incubated with two connector oligonucleotides together with DNA ligase for 30 min at 37°C. If the two secondary antibodies are in close proximity, this step allows the connector oligonucleotides to hybridize to the PLA probes and form a circular DNA strand after ligation. Incubation, for 100 min at 37°C, with DNA polymerase consequently leads to rolling circle amplification (RCA), the products of which are detected using fluorescently-labeled complementary oligonucleotides. Cells were washed with Duolink In Situ wash buffers following each of these steps. In the last wash, 1 μg/ml DAPI was added to the cells for 5 min at room temperature in the dark to stain the nuclei and coverslips were set in Mowiol (polyvinyl alcohol 4-88, Fluka), on glass slides. Cells were analyzed using a confocal Leica SP5-SMD microscope, with a LEICA 63x/1.4 OIL HCX PL APO CS objectives. Images were acquired by the Confocal head TCS SP5 II using the Leica Application Suite X software. Images were processed using Image J. When indicated, z-stacks of images were submitted to maximum intensity projection. The number of dots per transfected cell was estimated in one slice, in around 100 cells in each condition, with an automated procedure using plugins from the Image J software.

### Immunofluorescence

BHK and Neuro2Acells were seeded onto glass coverslips. Primary CGNs were cultured on coverslips previously coated with laminin (16,67 μg/ml) and poly-D-lysine (33,3 μg/ml). Cells and neurons were treated as described in the figure legends and then fixed with 4% paraformaldehyde. Overexpressed HA-NFATc3 and endogenous Trim39 were detected using anti-HA (1:500) and anti-Trim39 (from Proteintech 1:200, from Origene 1:400) antibodies respectively, as described previously (Iréna Lassot et al., 2018). GFP and mCherry-fused proteins were visualized by fluorescence and nuclei were stained with DAPI. Coverslips were analyzed by conventional or confocal microscopy, as mentioned in the figure legends. Image acquisition and analysis were performed on work stations of the Montpellier imaging facility (Leica DM600 for conventional microscopy, Leica SP5-SMD for confocal microscopy). For quantification of NFATc3 nuclear localization, BHK cells with predominant cytoplasmic or nuclear localization of NFATc3 were counted, in a blinded manner, among double GFP/HA positive cells. At least 100 double positive cells were scored for each experiment and each condition.

### In vivo ubiquitination of NFATc3

Neuro2A or BHK cells cultured in 60 mm or 100 mm dishes were transfected with pCI-HA-NFATc3 together with a plasmid expressing eight His_6_-tagged ubiquitin (His-Ub), or empty pCI, in the presence or the absence of pCI-Flag-Trim39, pCI-Flag-Trim39-ΔRING, pCI-Trim17-GFP or a combination of these plasmids. Following 24 h transfection, the medium was supplemented with 20 μM MG-132 for 6 h. Then, cells were harvested in 1 ml PBS without Ca^2+^ and Mg^2+^ supplemented with 10 μM MG-132. In some experiments, a 100 μl sample of the homogenous cell suspension was taken for input analysis and transfection efficiency control. After centrifugation, the pellet from the remaining 900 μl cell suspension was homogenized in 1 ml lysis buffer B (6 M guanidinium-HCl, 0.1 M Na_2_HPO_4_/NaH_2_PO_4_, 10 mM Tris-HCl [pH 8.0]) supplemented with 5 mM imidazole, 510 mM β-mercaptoethanol, 0.5 M NaCl and 10 mM NEM. The lysate was sonicated, cleared by centrifugation at 1,500 × g for 5 min at room temperature. In some experiments, input analysis was made at this stage by precipitating 50 μl of the resulting supernatants with TCA. The remainder of each extract was added to 40 μl magnetic nickel beads (MagneHis™ Ni-Particles, Promega). Beads were rotated for 2 h at room temperature to purify ubiquitinated proteins and washed once with lysis buffer B supplemented with 20 mM imidazole, 0.5 M NaCl and 10 mM NEM, once with 8 M urea, 0.1 M Na_2_HPO_4_/NaH_2_PO_4_, 10 mM Tris-HCl (pH 8.0), 20 mM imidazole,0.5 M NaCl and 10 mM NEM, three times with 8 M urea, 0.1 M Na_2_HPO_4_/NaH_2_PO_4_, 10 mM Tris-HCl (pH 6.3), 20 mM imidazole,0.5 M NaCl, 10 mM NEM, 0.2% Triton X-100, once with 8 M urea, 0.1 M Na_2_HPO_4_/NaH_2_PO_4_, 10 mM Tris-HCl (pH 6.3), 20 mM imidazole,0.5 M NaCl, 10 mM NEM, 0.1% Triton X-100 and once with 8 M urea, 0.1 M Na_2_HPO_4_/NaH_2_PO_4_, 10 mM Tris-HCl (pH 6.3), 10 mM imidazole, 10 mM NEM. Materials bound to the beads were eluted by the addition of 3 × Laemmli sample buffer and boiling for 5 min. These purification products and initial total lysates were resolved by SDS-PAGE and analyzed by immunoblotting.

### Production of recombinant TRIM proteins

BL21-CodonPlus®(DE3)-RP competent cells (Agilent) were transformed with the expression vectors pGEX-4T1 expressing GST-Trim39 fusion proteins (WT and mutants). Protein expression was induced by the addition of 500 μM IPTG and was carried out at 20°C overnight in the presence of 100 μM ZnCl_2_ and 200 μM MgSO_4_. Bacteria were harvested by centrifugation and resuspended in BugBuster® Protein Extraction Reagent (Millipore #70584-4) supplemented with protease inhibitor cocktail (cOmplete EDTA-free, Sigma-Aldrich). Bacterial suspensions were incubated for 30 min at room temperature with 1 mg/ml lysozyme (Fluka) and further lysed by sonication. The soluble protein fraction was recovered by centrifugation. GST fusion proteins were isolated by binding to glutathione magnetic beads (MagneGST™ Glutathione Particles, Promega) for 30 min at room temperature on a rotating wheel. The beads were then washed three times with 0.65 M NaCl and three times with PBS.

ArcticExpress (DE3) competent cells (Agilent) were transformed with the expression vector pET-15-HIS-TRIM39 and pMAL-c2-TRIM17 (expressing MBP-TRIM17 fusion protein). Bacteria were grown overnight in LB medium supplemented with Ampicillin and Gentamycin (20 μg/ml). Recombinant protein expression was induced by the addition of 1 mM IPTG and was carried out at 12°C overnight in the presence of 100 μM ZnCl_2_ and 200 μM MgSO_4_. Bacteria were harvested by centrifugation. Pellets were resuspended in bacterial lysis buffer (50 mM Tris-HCl [pH 8,6], 0.5 M NaCl, 50 mM MgSO_4_) supplemented with lysozyme and protease inhibitors, and frozen in liquid nitrogen to lyse bacteria. Lysates were then cleared by centrifugation. MBP-TRIM17 proteins were purified on amylose resin (New England BioLabs, #E8021L) and then eluted in a buffer containing 20 mM maltose before dialysis in PBS. HIS-TRIM39 proteins were purified on Ni-NTA agarose beads (Qiagen, #1018244) and then eluted in a buffer containing 0.5 M imidazole before dialysis in PBS.

### *In vitro* ubiquitination assay

NFATc3 was first transcribed and translated *in vitro*. For this, 1 μg of pCS2-HA-NFATc3 was incubated for 2 h at 30°C in 50 μl of the TNT® SP6 coupled wheat germ extract system (Promega, #L5030), according to the instructions of the manufacturer. For each ubiquitination condition, the equivalent of 3 μl of the *in vitro* translation reaction was immunopurified using 1 μl rat anti-HA antibody and 10 μl protein G-agarose beads in a buffer containing 50 mM Tris-HCl (pH 7.5) and 50 mM NaCl buffer for 1 h. Beads were washed 3 times in the same buffer, as described above for co-immunoprecipitation. Then, beads carrying NFATc3 were incubated in 20 μl of ubiquitination reaction buffer (50 mM Tris-HCl [pH 7.5], 50 mM NaCl, 4 mM ATP, 4 mM MgCl2, 2 mM DTT, 10 mM phosphocreatine, 0.5 U creatine kinase, 20 μM ZnCl_2_), in the presence of 50 ng human recombinant His-tagged ubiquitin-activating enzyme E1 (from BostonBiochem, #E-304), 500 ng human recombinant His-tagged ubiquitin-conjugating enzyme (E2) Ube2d3 (from BIOMOL International, #U0880), in the presence or the absence of 10 μg N-terminal-His-tagged ubiquitin (Sigma-Aldrich, #U5507), and ≈ 2 μg of purified recombinant mouse GST-Trim39 (WT) or GST-Trim39-C49S/C52S, or ≈ 2 μg of purified recombinant human His-TRIM39 in the presence or the absence of ≈ 2 μg MBP-TRIM17. Reactions were incubated at 37°C for 1 h. Beads were washed once and the reaction was stopped by adding 10 μl of 3 × Laemmli sample buffer and by heating at 95°C for 5 min. The samples were analyzed by SDS-PAGE and immunoblotting with anti-NFATc3, anti-Trim39 and anti-TRIM17 antibodies.

### *In vitro* SUMOylation assay

WT NFATc3 and its KallR and EallA mutants were first transcribed and translated *in vitro* as described above. Equivalent amounts of the different forms of NFATc3 (between 1,5 and 6 μl of the *in vitro* translation reaction) were incubated for 1 h 30 min at 37°C in the presence of 3 μg recombinant SUMO1, 150 ng recombinant His-tagged Aos1/Uba2 (E1 enzyme), 100 ng recombinant Ubc9 (E2 enzyme) and 300 ng recombinant GST-PIASxα (E3 enzyme) in 20 μl shift-assay buffer (20 mM Hepes [pH 7.3], 110 mM KOAc, 2 mM Mg(OAc)_2_, 0.5 mM EGTA, 1 mM DTT, 0.05% Tween 20, 0.2 mg/ml ovalbumin, 1 μg/ml leupeptin, 1 μg/ml aprotinin, 1 μg/ml pepstatin) supplemented with 1 mM ATP. Recombinant proteins used in this assay were produced and purified as previously described (Bossis et al., 2005). Negative controls (time 0) were obtained by mixing all reagents directly into loading buffer. Reaction products were separated by SDS-PAGE (Tris-acetate gels), transferred to PVDF membranes and analyzed by western blot using anti-NFATc3 antibody.

### RNA preparation and quantitative RT-PCR

Total RNA was extracted using the RNeasy kit (Qiagen) and treated with the DNase I from the DNA-free™ kit (Thermo Fisher Scientific) according to manufacturer’s instructions. RNA was used to perform a two-step reverse-transcription polymerase chain reaction (RT-PCR). In brief 1 μg of total RNA was reverse-transcribed using 200 U reverse transcriptase Superscript II (Thermo Fisher Scientific) in the presence of 2.5 μM N6 random primers and 0.5 mM dNTP. The equivalent of 6 ng of resulting cDNA was used as a template for real time PCR using a Mx3000P thermocycler (Agilent) with a home-made SYBR Green qPCR master mix (Lutfalla & Uze, 2006). PCR reactions were performed in 10 μl in the presence of 200 nM primers. Thermal cycling parameters were 10 min at 95°C, followed by 40 cycles of 95°C for 30 s, 64°C for 30 s and 72°C for 30 s. Specific primers used to amplify mouse *Trim39* and mouse *Trim17* are listed in supplementary Table 1. Data were analysed and relative amounts of specifically amplified cDNA were calculated with MxPro software (Agilent), using the mouse *Gapdh* amplicon as a reference.

### Protein sequence analysis

The sequence of mouse Trim39 (GenBank: NM_024468) was analyzed by using the prediction web-based tools JASSA (Joined Advanced SUMOylation site and SIM analyzer, (http://www.jassa.fr/) and GPS SUMO (group-based prediction system, http://sumosp.biocuckoo.org/online.php) to identify putative SUMO-interacting motifs.

### Production of SUMO chains and GST-pull down

Recombinant free SUMO-2 and poly-SUMO-2 chains were produced in bacteria co-expressing His-SUMO-2, Aos1/Uba2 (SUMO conjugating E1 enzyme) and Ubc9 (SUMO E2 conjugating enzyme). For this, BL21 competent cells were transformed with the plasmid pE1-E2-His-Su2 (described in (Brockly et al., 2016)). The transformed bacteria were grown with strong agitation (210 rpm) at 37°C until the OD reaches 0.4-0.6. Protein expression was induced by adding 1 mM IPTG for 6 h at 25°C. The bacteria were resuspended in 30 ml of bacterial lysis buffer, frozen in liquid nitrogen and stored at −80°C. The resuspended bacteria were thawed and supplemented with lyzozyme (1 mg/ml), 8 mM ß-mercaptoethanol, 1 μg/ml aprotinin, 1 μg/ml leupeptin, 1 μg/ml pepstatin and incubated for 1 h on ice before centrifugation at 100,000 × g for 1 h at 4°C. The supernatant was loaded on a Ni-NTA column equilibrated in bacterial lysis buffer supplemented with 8 mM ß-mercaptoethanol, 0.5% Triton X-100, 10 mM imidazole and protease inhibitors. The column was washed 3 times with 10 ml of the same buffer and eluted with 15 ml of Ni-NTA elution buffer (bacterial lysis buffer supplemented with 250 mM imidazole and protease inhibitors).

For GST-pull down, GST-Trim39 and its different SIM mutants were produced in bacteria and purified as described above. The quantity and the quality of the different forms of GST-Trim39 bound to glutathione magnetic beads was first estimated on a poly-acrylamide gel using Coomassie blue staining, by reference to known amounts of BSA. Beads binding approximately 1 μg of each form of GST-Trim39 were incubated with ≈ 1 μg SUMO-2 chains for 1 h at room temperature in 200 μl shift assay buffer (20 mM Hepes [pH 7.3], 110 mM KOAc, 2 mM Mg(OAc)_2_, 0.5 mM EGTA, 1 mM DTT, 0.05% Tween 20, 0.2 mg/ml ovalbumin, 1 μg/ml leupeptin, 1 μg/ml aprotinin, 1 μg/ml pepstatin). Beads were recovered by centrifugation and washed 4 times with PBS. Material bound to the beads was eluted by the addition of 3 × Laemmli sample buffer and incubation at 95°C for 5 min. Both GST-Trim39 proteins and bound SUMO-2 chains were analyzed by immunoblotting.

### Assessment of neuronal apoptosis

After 6 days *in vitro* (DIV), transfected or transduced CGNs were maintained in initial culture medium (control) or were washed once and incubated in serum-free BME supplemented with L-Gln, HEPES, antibiotics and 1 μM (+)-MK-801, and containing 5 mM KCl (K5 medium) for the indicated times. Neurons were then stained with DAPI and mounted on glass slides in Mowiol. In experiments in which the CGNs were transfected with GFP, GFP-NFATc3 or GFP-NFATc3-EallA, apoptosis was assessed among GFP-positive neurons, by examining neuronal morphology and nuclear condensation. For each experiment and each condition, at least 100 GFP-positive neurons were scored in a blinded manner. In experiments in which CGNs were transduced with shRNA-expressing lentiviruses, apoptosis was estimated by counting the percentage of condensed nuclei in five random fields for each condition (more than 500 cells).

### Statistics

Statistical analyses were performed using GraphPad Prism version 7.0c for Mac OS X (GraphPad Software, San Diego California USA, www.graphpad.com). Unless stated, data are representative of at least three independent experiments.

## Acknowledgements

This work was supported by the Centre National de la Recherche Scientifique (CNRS), the Institut National de la Santé et de la Recherche Médicale (INSERM), the Université de Montpellier, La Fondation de l’Association pour la Recherche contre le Cancer (ARC), La Ligue contre le Cancer and La Fondation pour la Recherche Médicale (FRM). This article is based upon work from COST Action (PROTEOSTASIS BM1307), supported by COST (European Cooperation in Science and Technology). We would like to thank the staff of the Montpellier Genomic Collection platform for providing human TRIM39 and human TRIM17 cDNA clones. We acknowledge the imaging facility MRI (Montpellier Ressources Imagerie), member of the national infrastructure France-BioImaging infrastructure supported by the French National Research Agency (ANR-10-INBS-04, «Investments for the future». We are grateful to Frédérique Brockly for the production and purification of recombinant proteins. We thank Drs Dimitris Liakopoulos and Manuel Rodriguez for interesting discussions.

