## supplemental figures and table for "Trim39 regulates neuronal apoptosis by acting as a SUMO-targeted E3 ubiquitin-ligase for the transcription factor NFATc3"

#### Figure S1

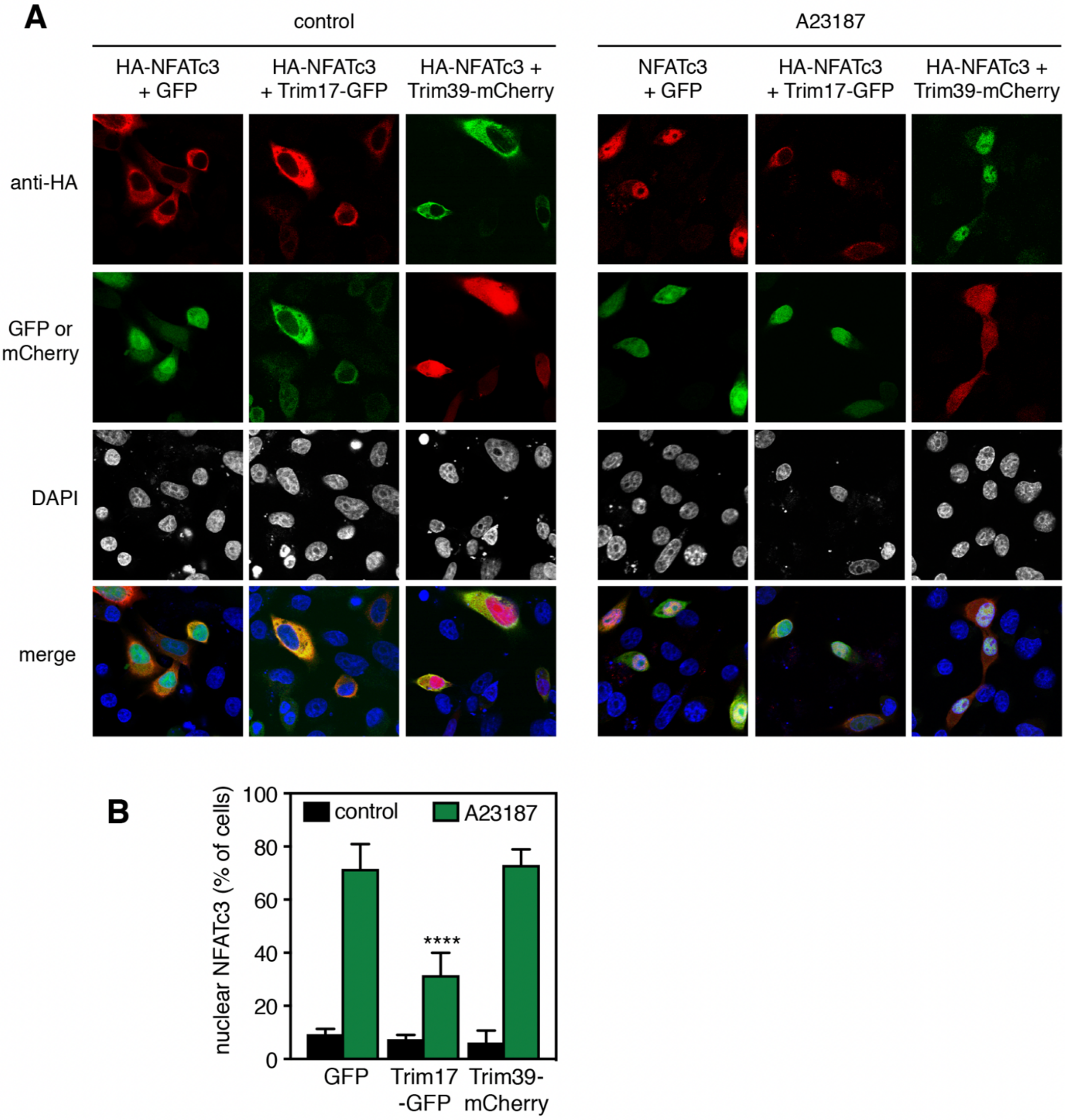

**Figure S1. Trim39 does not alter calcium-induced nuclear translocation of NFATc3. A.** BHK cells were transfected with HA-NFATc3 together with GFP (negative control) or Trim17-GFP, or Trim39-mCherry for 24 h. Then, cells were left untreated (control) or deprived of serum for 3h and treated with 1 µM of the calcium ionophore A23187 in serum-free medium for an additional 30 min before fixation. NFATc3 was detected using an anti-HA antibody and visualized by confocal microscopy. GFP, Trim17-GFP and Trim39-mCherry were visualized by GFP or mCherry fluorescence and nuclei were stained with DAPI. **B.** Quantification of the nuclear localization of NFATc3 in experiments conducted as in A. The percentage of cells showing NFATc3 mainly in the nucleus was determined among the population of cells expressing both HA-NFATc3 and either GFP, Trim17-GFP or Trim39-mCherry. Data are the means ± SD of three independent experiments. **** $P<0.0001$ significantly different from the corresponding value obtained in cells transfected with GFP and NFATc3 (two way ANOVA followed by Dunnett's multiple comparisons test).

### Figure S2

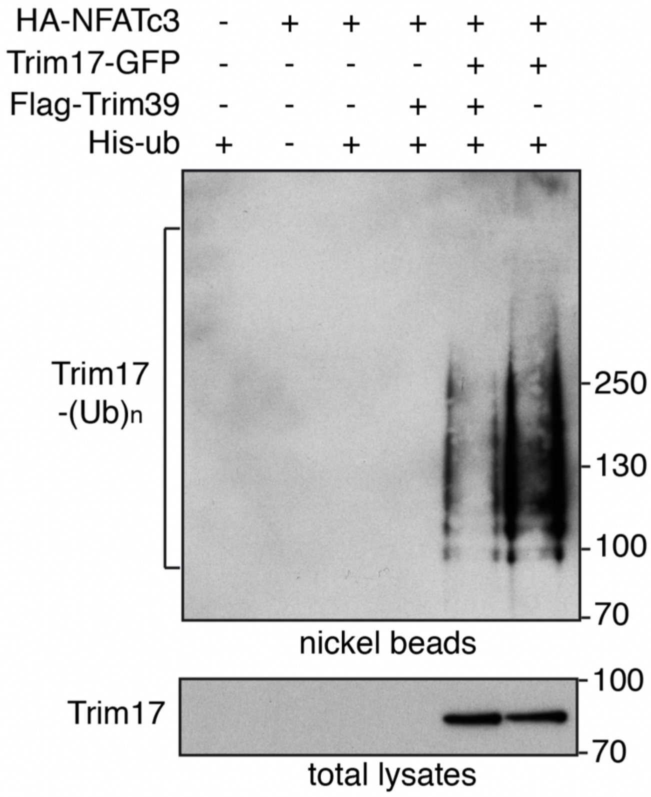


**Figure S2. Trim39 decreases the ubiquitination of Trim17.** The PVDF membrane presented in Figure 4A was stripped and blotted with an anti-GFP antibody.

### Figure S3

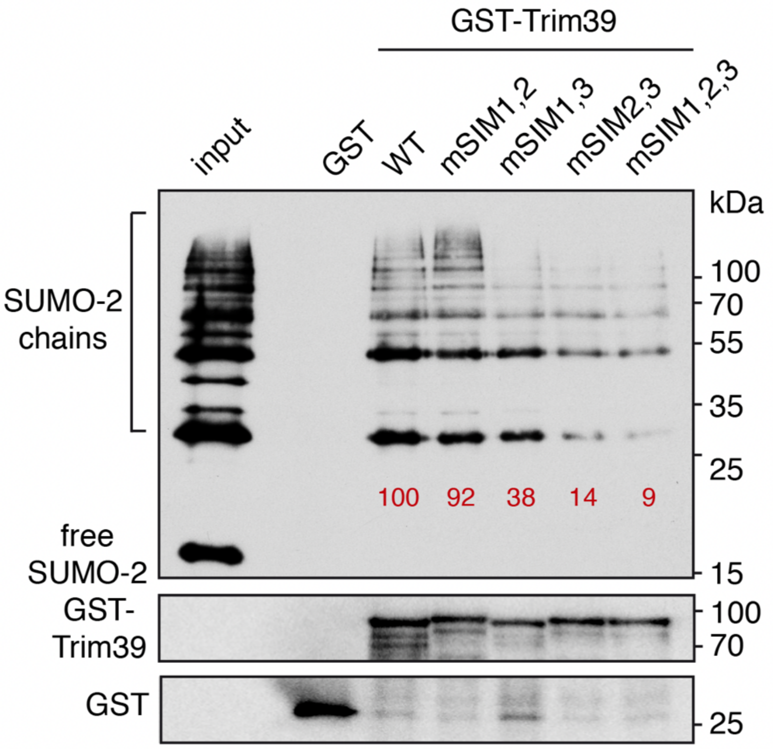


**Figure S3. SIM3 is the predominant SIM involved in the interaction of Trim39 with SUMO-2 chains.** Recombinant GST, GST-Trim39 and its different double or triple SIM mutants were purified using glutathione beads and subsequently incubated with purified recombinant SUMO-2 and SUMO-2 chains. Material bound to the beads was eluted and analyzed by western blot using anti-SUMO and anti-GST antibodies. A small fraction of the SUMO-2 chains was also loaded on the gel (input) for comparison. The intensity of bound SUMO-chain bands was quantified and normalized by the intensity of corresponding GST-Trim39 bands. Relative values are indicated in red.

**Supplementary Table 1: list of the primers used to generate described constructs**

| PCR primer pairs used for constructs: | | |
|---|---|---|
| Construct | Forward (5' – 3') | Reverse (5' – 3') |
| NFATc3-E437A | GAATTGAAAATAGCAGTGCAACCTAAAAC | GTTTTAGGTTGCACTGCTATTTTCAATTC |
| NFATc3-E706A | GATGAAGCAAGCACAAAGAGAAGAC | GTCTTCTCTTTGTGCTTGCTTCATC |
| NFATc3-E1015 | CATTAAACCTGCACCTGAAGATCAAG | CTTGATCTTCAGGTGCAGGTTTAATG |
| Trim39 | ATAGAATTCATGGCAGAGACAAGTCTG | TTATCTAGATCATTCCCAATCTGTTGG |
| Trim39-ΔRING | CGAGAATTCTCCCGATACCGG | CGACTCGAGTCATTCCCAATC |
| Trim39-mSIM1 | GAGTATCTGAAGGAGCCAGCTGCTGCTGAATGTGGGCACAAC | GTTGTGCCCACATTCAGCAGCAGCTGGCTCCTTCAGATACTC |
| Trim39-mSIM2 | GAAGACCAGGAGGCTGCATGTGCCGCCTGTGCAATTTCTCATACCC | GGGTATGAGAAATTGCACAGGCGGCACATGCAGCCTCCTGGTCTTC |
| Trim39-mSIM3 | CGAAGAGCAGCAGACAGCAGCAGCGCGAGCAGAGGAAGAAGAACAG | CTGTTCTTCTTCCTCTGCTGCCGCTGCTGCTGTCTGCTGCTCTTCG |
| Trim39-mCherry 1st PCR | ATAGAATTCATGGCAGAGACAAGTCTG | CTCCTCGCCCTTGCTCACTTCCCAATCTGTTGGG |
| Trim39-mCherry 2nd PCR | CCCAACAGATTGGGAAGTGAGCAAGGGCGAGGAG | ATATCTAGATTATCTAGATCCGGTGGATC |
| Trim39-mCherry 3rd PCR | ATAGAATTCATGGCAGAGACAAGTCTG | ATATCTAGATTATCTAGATCCGGTGGATC |
| GST-Trim39 | CGAGAATTCGCAGAGACAAGTC | CGACTCGAGTCATTCCCAATC |
| Trim39-C49S/C52S | GGCACAACTTCTCCAAAGCGTCCATTACCGCGTGG | CCAGCGGGTAATGGACGCTTTGGAGAAGTTGTGCC |
| TRIM17 | CGAGAATTCGAGGCTGTGGAACTCGCC | CGAGTCGACCTATCCTTTCACCCACATGGTCAC |
| Primers used for RT-qPCR | | |
| cDNA | Forward (5' – 3') | Reverse (5' – 3') |
| Trim39 | TGGAGGTGACTTCAGTATCCAT | TCACATCCGCAATTAGCTGTT |
| Trim17 | ACTGAGTGGCAGGAGAGAGTGAA | CCAGAAACAAGACCACCTTCTGA |
| Gapdh | AAATGGTGAAGGTCGGTGTG | CCTTGACTGTGCCGTTGAAT |
